# Deciphering abdominal aortic diseases through T-cell clonal repertoire of perivascular adipose tissue

**DOI:** 10.1101/2023.12.05.570098

**Authors:** Luca Piacentini, Chiara Vavassori, Pablo Werba, Claudio Saccu, Rita Spirito, Gualtiero I. Colombo

## Abstract

Recent studies suggested that immune-mediated inflammation of perivascular adipose tissue (PVAT) of abdominal aortic aneurysms (AAA) contributes to disease development and progression. Whether the PVAT of AAA is characterized by a specific adaptive immune signature remains unknown. To investigate this hypothesis, we sequenced the T-cell receptor β-chain (TCRβ) in the PVAT of AAA patients and compared with patients with aortic occlusive disease (AOD), who share with the former anatomical site of the lesion, risk factors but differ in pathogenic mechanisms. Our results demonstrate that AAA patients have a lower repertoire diversity than those with AOD and significant differences in V/J gene segment usage. Furthermore, we identified a set of 7 public TCRβ clonotypes that distinguished AAA and AOD with very high accuracy. We also found that the TCRβ repertoire differentially characterizes small and large AAA (aortic diameter <55 mm and ≥55 mm, respectively). This work supports the hypothesis that T-cell-mediated immunity is fundamental in AAA pathogenesis and opens up new clinical perspectives.

**Summary:** Different immune mechanisms may play a key role in the pathogenesis of distal aortic aneurysm and aortoiliac occlusive disease. The TCRβ repertoire of perivascular adipose tissue differs between the two pathologic conditions, suggesting the involvement of specific antigen-specific immune responses.

## Introduction

Abdominal aortic aneurysm (AAA) is a chronic, life-threatening vascular disease for which no early detection markers and pharmacological therapy to stop or delay AAA progression are available to date, making it a serious and challenging health concern. Ineffective pharmacological therapy stems from an inadequate understanding of the etiopathogenesis of AAA (Kent et al., 2010; Golledge, 2019). The only treatment option is surgical repair or stenting to prevent vessel rupture. Understanding the pathogenetic mechanisms of AAA is therefore crucial to finding new effective therapeutic solutions.

Perivascular adipose tissue (PVAT) and its immune cell infiltrates are recognized to play a central role in vascular homeostasis and in the pathogenesis of large vessel diseases, including atherosclerotic and aneurysmal lesions (Queiroz and Sena, 2020; Guzik et al., 2017; Huang Cao et al., 2017). Immune-inflammatory mechanisms engaging PVAT have been recently identified in patients with AAA and appear to be involved in the development and progression of this pathology (Piacentini et al., 2019, 2020a, 2022; Folkesson et al., 2017). The emergence of an antigen-specific T-cell immune response, *e*.*g*. activation and proliferation of T lymphocytes, and/or its ineffective control has been hypothesized to trigger a self-inflammatory loop that leads to the growth of AAA (Piacentini et al., 2019; Sagan et al., 2019; Li et al., 2022; Zhou et al., 2015; Suh et al., 2020).

T-cells are an integral component of the adaptive immune response and a predominant immune cell population in PVAT of AAA (Sagan et al., 2019). Interacting with antigens presented by human leukocyte antigens (HLA) molecules on antigen-presenting cells (APCs), T-cell receptors play an essential role in the activation and proliferation of T-cells themselves. The complementarity determining region 3 (CDR3) is the most relevant portion of the TCRβ whose high variability results from the countless combinations of the variable (V), diversity (D), and joining (J) gene segments (and palindromic, random nucleotide additions or deletions as well) and determines the most important part of the TCRβ with which the peptide antigen binds.

The relationship between adaptive immunity and AAA is incompletely understood and whether a definite T-cell signature in PVAT can identify AAA is an open question that has not been addressed so far. The identification of specific hallmarks of the clonal repertoire of TCRβ and distinctive TCRβ CDR3 amino acid sequences would provide decisive evidence for the involvement of adaptive immune response in the pathogenesis of AAA.

In this study, we tested the hypothesis that a distinctive repertoire of T lymphocytes characterizes and marks the PVAT in AAA patients.

To this aim, we sought to identify and quantify the clonal repertoire of T lymphocytes (or T-cells) by deep sequencing of the T-cell receptor β-chain (TCRβ) in the PVAT (of both lesion sites and non-lesioned segments) of AAA patients. As a meaningful comparison, we analyzed TCRβ repertoires from patients with abdominal aortic occlusive disease (AOD), due to the common anatomical site of the lesions and shared risk factors but pathogenetic differences (Biros et al., 2015; Criqui and Aboyans, 2015; Horimatsu et al., 2017; Piacentini et al., 2020b).

The analysis of the TCR repertoire allows us to characterize the immune status of host T-cells in different morbid conditions (Chowdhury et al., 2022; Tang et al., 2019). Quantitative assessment of T-cell clones can be of pathogenic and clinical importance (Farmanbar et al., 2019; Mitchell and Michels, 2020).

## Results

### TCRβ repertoire diversity and overlap analysis of AAA and AOD

We estimated the diversity of the TCRβ repertoire in PVAT from the lesion site of AAA patients (n=28) and compared it with that in the PVAT of the lesion site of the AOD group (n=11). Both AAA and AOD patients showed some heterogeneity in the extent of diversity, although AAA patients had a significantly restricted clonal repertoire, which was 67% smaller than in AOD patients (**Figure 1A**). The smaller clonal repertoire width in the PVAT of AAA patients reflects a reduced diversity of the repertoire suggesting an antigen-specific immune response mechanism. Repertoire clustering analysis revealed the presence of rather homogeneous groups of disease-related samples: in particular, the AAA samples clustered into three rather distinct groups, while most of the AOD patients clustered together (**Figure 1B**). The precise grouping of different types of technical replicates when included in a clustering analysis, as described in the Methods section, gives confidence in the result obtained (**Supplementary Figure S1**).

**Figure 1.**
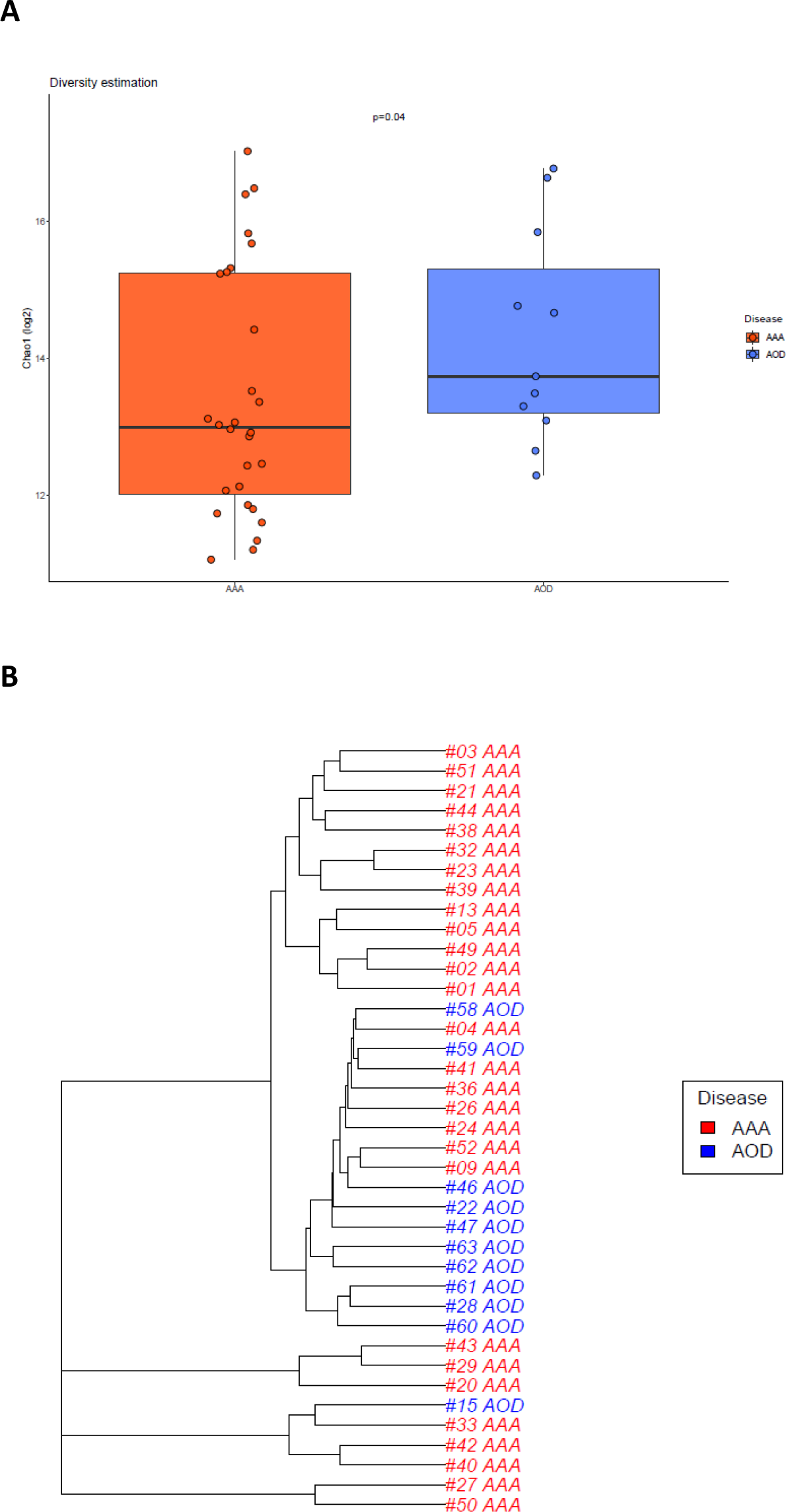
TCRβ repertoire diversity and overlapping in AAA and AOD. Panel (**A**) shows the box plot of the Chao-1 diversity estimate in AAA (red) and AOD (blue) patients. The one-tailed Wilcoxon test was used to assess differences between the two groups. Significance threshold was set at P-value<0.05. Panel (**B**) shows the hierarchical clustering of the clonotype similarity measure, calculated as the clonotype sum of geometric mean frequencies (F2; see Methods section). Clonotypes were defined as those with identical CDR3 amino acid sequence and V segment.

### Homeostatic clonal space in AAA and AOD

We evaluated the proportions of rare to hyperexpanded clonotypes in each patient of the two disease groups. Hyperexpanded clonotypes occurred in almost all the PVAT of AAA and AOD patients, despite having a private characteristic (**Figure 2A;** *cf*. Methods section for definition of ‘private’ and ‘public’ clonotypes.). The relative abundance of hyperexpanded clonotypes did not differ significantly between the two groups. In contrast, we found that the frequency of large and medium clonotypes was higher in AAA than in AOD (ratio AAA *vs*. AOD= 1.7 and 1.3, respectively), whereas the frequency of small and rare clonotypes was concordantly lower in AAA patients (ratio AAA *vs*. AOD= -1.75 and -1.8, respectively), although not statistically significant (**Figure 2B**). Yet, the frequency of rare or small clonotypes showed a significant positive correlation (ρ=0.98 and ρ=0.93, respectively, with p<1x10^−10^) with the Chao-1 diversity estimate, whereas the frequency of medium, large, or hyperexpanded clonotypes showed a lower but still significant negative correlation (ρ=-0.78, ρ=-0.82 and ρ=-0.66, respectively, with p<1x10^−5^ for all) with the Chao-1 diversity estimate, suggesting that the clonal diversity in these samples is mainly driven by the fraction of rare and small clonotypes (**Supplementary Figure S2**). In addition, the occurrence of predominantly private hyperexpanded clonotypes in patients with AAA and AOD may have led to a less clear-cut overlap of repertoires in each disease group, as shown by the clustering analysis. Indeed, the calculation of the clonotype sum similarity measure of geometric mean frequencies (F2), used in our overlap analysis (see Methods section), limited the effect of technical bias (*e*.*g*., cross-sample contamination), but at the same time penalizes the similarity score of two samples with highly-frequency private clonotypes, which are the hyperexpanded clonotypes.

**Figure 2.**
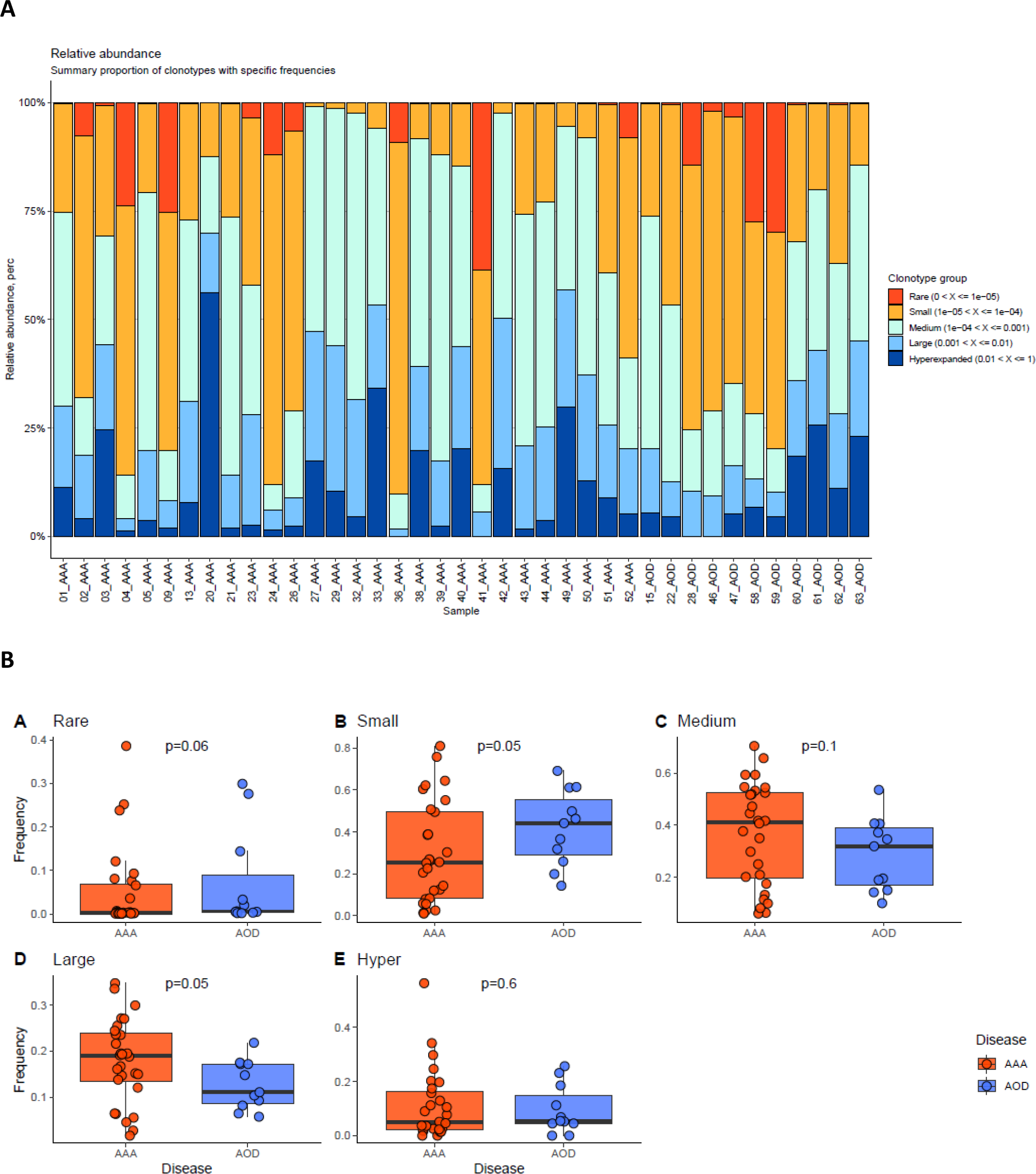
Clonal space homeostasis in AAA and AOD. Panel (**A**) displays the relative abundance of the cumulative frequency of clonotypes defined as Rare ≤0.00001; 0.00001<Small≤0.0001; 0.0001<Medium≤0.001; 0.001<Large≤0.01; and Hyperexpanded > 0.01 for each sample. Panel (**B**) shows the box plots of clonotypes frequencies in the two disease groups. The one-tailed Wilcoxon test was used to assess differences between AAA (red) and AOD (blue) patients. The significance level was set at P<0.05.

### Differential V and J Segment Usage between AAA and AOD

We tested the differential usage for all the V and J gene segments expressed in AAA *vs*. AOD samples. We found that the frequency of TRBV20-1, TRBJ2-4, TRBJ1-3, and TRBJ2-7, showed a significant difference between AAA and AOD (**Figure 3**), suggesting that the two patient groups are characterized by a preferential selection of clonotypes with specific V/J segments (also known as TCR bias). **Supplementary Data 1a** and **Supplementary Figures S3** summarize the results and statistics of the entire analysis.

**Figure 3.**
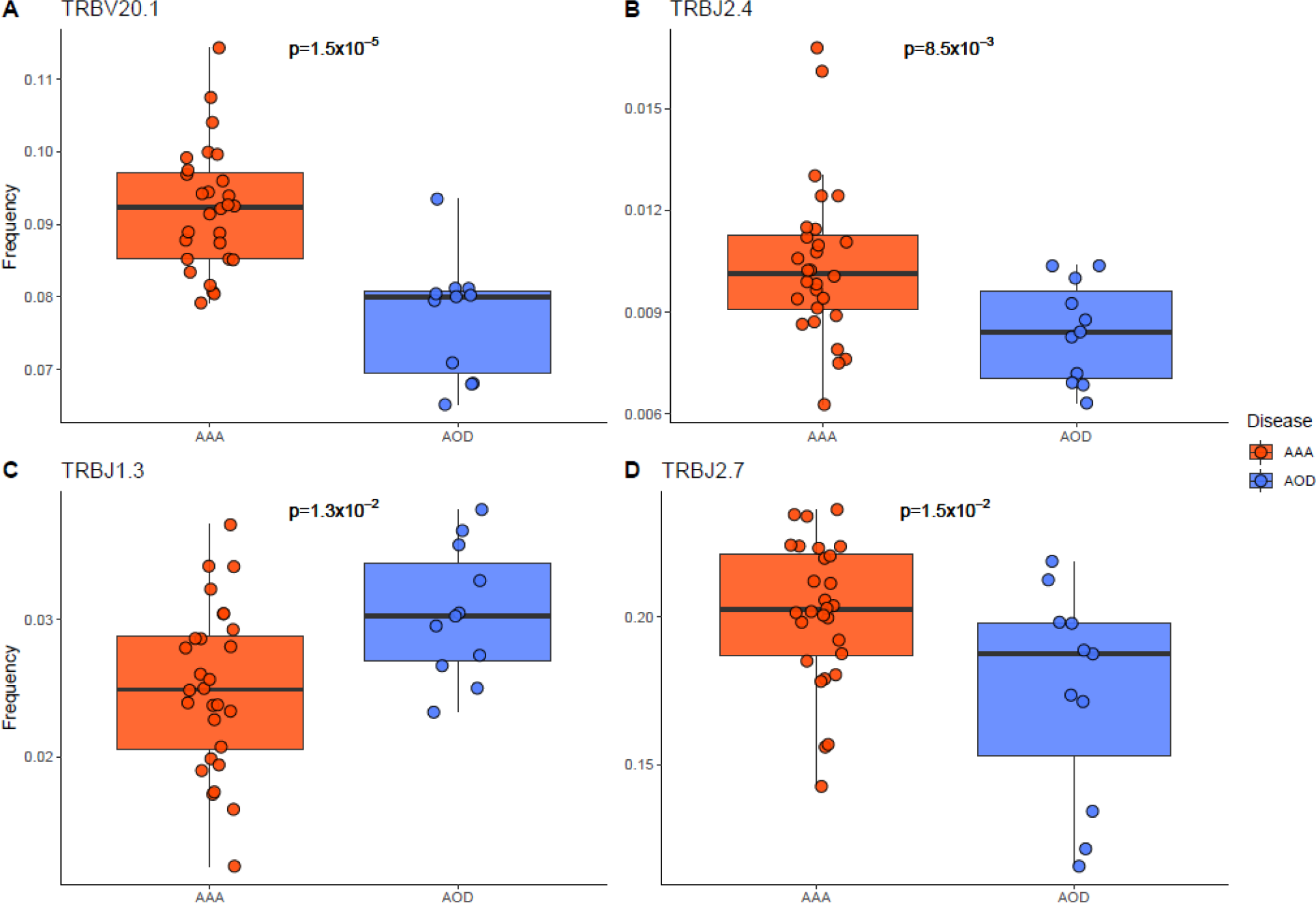
V/J segment usage differences between AAA and AOD. Box plots show the distribution and significant differences of V and J segment frequencies in AAA (red) and AOD (blue) patients. The two-tailed Wilcoxon test was used to assess differences between the two groups, with significance set at P-value<0.05.

### Disease-specific TCRβ clonotypes

We further investigated the possibility to identify TCRβ clonotypes specifically associated with a disease based on their CDR3 amino acid sequences and V-segment (aaV). We initially identified 558,506 and 301,224 aaV clonotypes in AAA and AOD patients, respectively. The majority of these are private clonotypes (96% and 92% in AAA and AOD, respectively), whereas the number of public clonotypes, *i*.*e*. shared by at least 2 subjects regardless of disease group, was 22,813. Of these, 12,094 (53%) are shared between the two disease groups in at least one subject per group, whereas 8,425 (37%) and 2,294 (10%) are shared only within AAA and AOD patients, respectively. The 3.7-fold difference in the number of shared clonotypes between AAA and AOD can be attributed to the higher number of AAA patients in our study, which would result in the higher number of unique clonotypes observed in AAA. On the other hand, when we searched for public aaV clonotypes in a similar proportion of both AAA and AOD patients, *i*.*e*. 18% (the proportion of AAA patients that met the minimum sharing allowed in the AOD group), we still found uniquely aaV clonotypes in AAA, although their number was reduced to 29. Although the total number of disease-specific aaV may vary considerably depending on the number of subjects involved, it is noteworthy that a relevant proportion (almost one-fifth) of patients in each disease group share disease-specific clonotypes.

We then explored the differences in the proportion of all public clonotypes in AAA *vs*. AOD, regardless of whether the clonotypes were shared within or between the two disease groups. We found 115 unique clonotypes that were differentially distributed between AAA and AOD (p<0.05; **Supplementary Data 1b**). From these, using a recursive feature elimination procedure and a machine learning approach, we identified a signature of 7 public clonotypes that allowed us to discriminate AAA from AOD patients with very high accuracy (AUC-ROC=0.99; sensitivity=0.99; specificity=0.92). We also found that this signature was restricted to the lesion site: in fact, only 4 out of 7 clonotypes were detected in the non-lesioned segment of the abdominal aorta PVAT, and prediction based on these 4 clonotypes did not effective discriminate AAA from AOD (AUC-ROC=0.68; sensitivity=0.84; specificity=0.43). As the reduced number of predictors may have affected the ability to discriminate between AAA and AOD, we also tried to classify AAA and AOD samples using only those 4 clonotypes but detected at the level of lesioned PVAT. Notably, we still obtained a good predictive performance (AUC-ROC=0.94; sensitivity=0.96; specificity=0.81), confirming that the tissue-specific localization of this clonotype signature capable of discriminating AAA from AOD was at the level of PVAT around the lesion. **Table 1** summarizes the overall measures of performance of these three cross-validated prediction models (*cf*. Methods section).

**Table 1.**
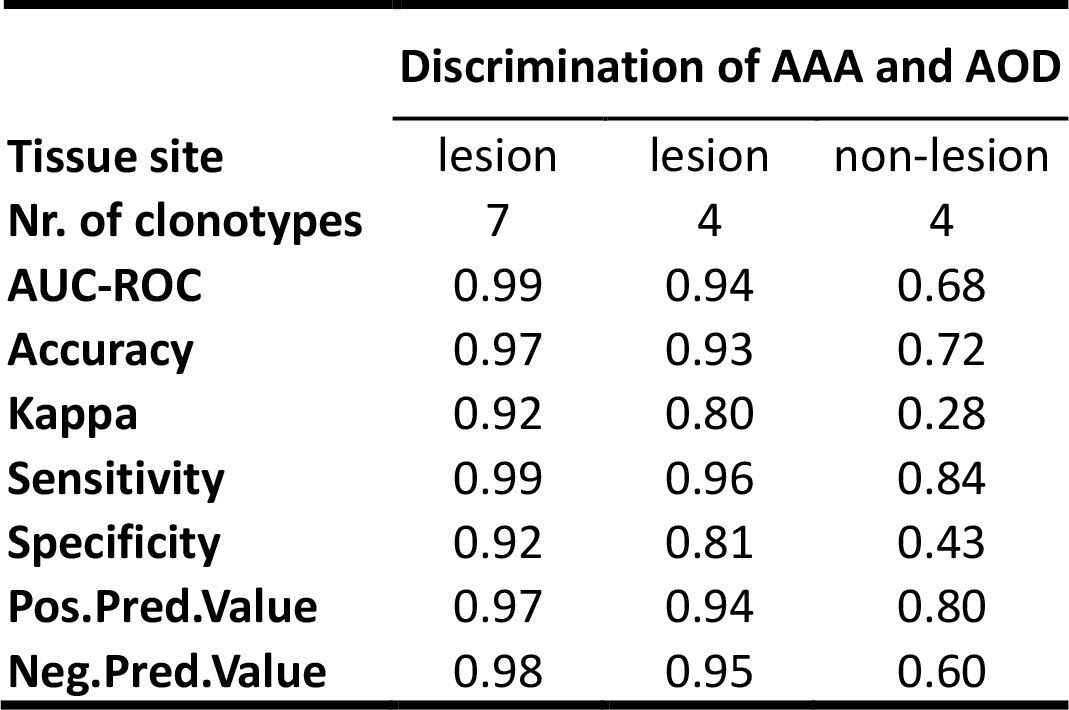
Discrimination of AAA from AOD through PVAT clonotypes. Summary of various performance measures of cross-validated models for predicting AAA and AOD PVAT samples using clonotype frequencies. Clonotypes were defined as those with identical CDR3 amino acid sequences and V segment. AUC, indicates the area under the curve; ROC, Receiver Operating Characteristic. Kappa is the Cohen’s Kappa.

|  | Discrimination of AAA and AOD |  |  |
| --- | --- | --- | --- |
| Tissue site | lesion | lesion | non-lesion |
| Nr. of clonotypes | 7 | 4 | 4 |
| AUC-ROC | 0.99 | 0.94 | 0.68 |
| Accuracy | 0.97 | 0.93 | 0.72 |
| Kappa | 0.92 | 0.80 | 0.28 |
| Sensitivity | 0.99 | 0.96 | 0.84 |
| Specificity | 0.92 | 0.81 | 0.43 |
| Pos.Pred.Value | 0.97 | 0.94 | 0.80 |
| Neg.Pred.Value | 0.98 | 0.95 | 0.60 |

### TCRβ repertoire differences between large and small AAA

In our previous work, we have shown that patients with different stages of AAA progression, *i*.*e*. small and large AAA, are characterized by different pathogenetic mechanisms (Piacentini et al., 2019, 2022). Thus, here we investigated whether the TCRβ repertoire might also differ in these two subgroups of AAA patients (see subgroup AAA patient characteristics in **Supplementary Table S1**).

Large *versus* small AAA did not show significant differences in either TCRβ repertoire diversity (**Supplementary Figure S4A**) or clonal homeostatic space (**Supplementary Figure S4B-F**). Instead, we found differences in the frequency of use of nine V gene segments, including TRBV3-1, TRBV6-2, TRBV4-3, TRBV10-2, TRBV5-1, TRBV3-2, TRBV6-1, TRBV7-2, and TRBV10-1 (**Figure 4**). No differences were found for the use of J gene segments (**Supplementary Data 1c)**.

**Figure 4.**
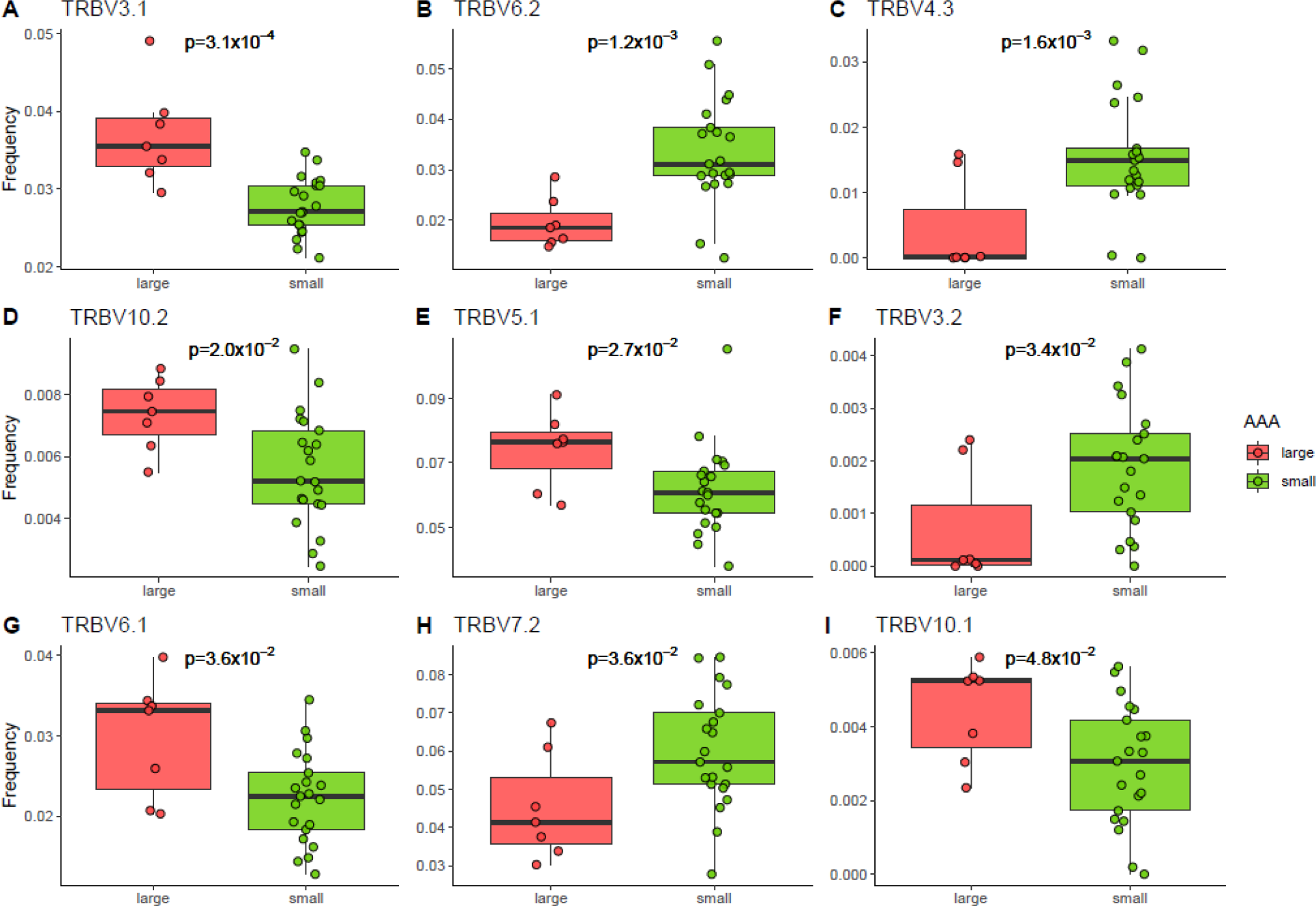
V segment usage differences between large and small AAA. Box plots show the distribution and significant differences of V segment frequencies in large (salmon) and small (green) AAA. The two-tailed Wilcoxon test was used to assess differences between the two groups, with significance set at P<0.05.

Applying the same approach as above, we also identified differences in the proportions of 9 public clonotypes (p<0.05; **Supplementary Data 1d**) in large *vs*. small AAA. Again, using a recursive feature elimination procedure over these differentially distributed clonotypes, we reduced the signature to 8 clonotypes that discriminated large from small AAA with high accuracy (AUC-ROC=0.99; sensitivity=0.88; specificity=0.97).

### Tissue-localized differences in the TCRβ repertoire in the PVAT surrounding the lesion compared with the non-lesioned area of the aorta in AAA and AOD

We also looked for possible specific tissue-localized differences of the TCRβ repertoire within each disease group. To this end, we performed a pairwise analysis of TCRβ repertoire of PVAT located at the lesion site *vs*. non-lesioned site on the subset of 24 and 10 paired-samples from AAA and AOD patients, respectively. The comparisons showed that there were no differences in TCRβ diversity and in clonal homeostatic space within either AAA or AOD patients (**Supplementary Figure S5**). Overlap and clustering analysis consistently showed that samples clustered per subject, suggesting that the private nature of the TCRβ repertoire within each subject prevailed over the overlap based on the disease type (**Supplementary Figure S6**). Instead, we observed a difference in V segment usage for TRBV3-1 and TRBV27 within AAA (**Figure 5A-B**) and for TRBV13, TRBV2, TRBV6-1, TRBV4-1 and TRBV7-6 within AOD samples (**Figure 5C-G**). For the J segments, no differences were found either within AAA or within AOD. Finally, we found only two aaV clonotypes that were differentially distributed in PVAT from the non-lesion site *vs*. the lesion site in AAA (p=0.04; sample frequency=0.25 [non-lesioned] *vs*. 0 [lesion]), whereas we did not find differentially distributed clonotypes within AOD patients.

**Figure 5.**
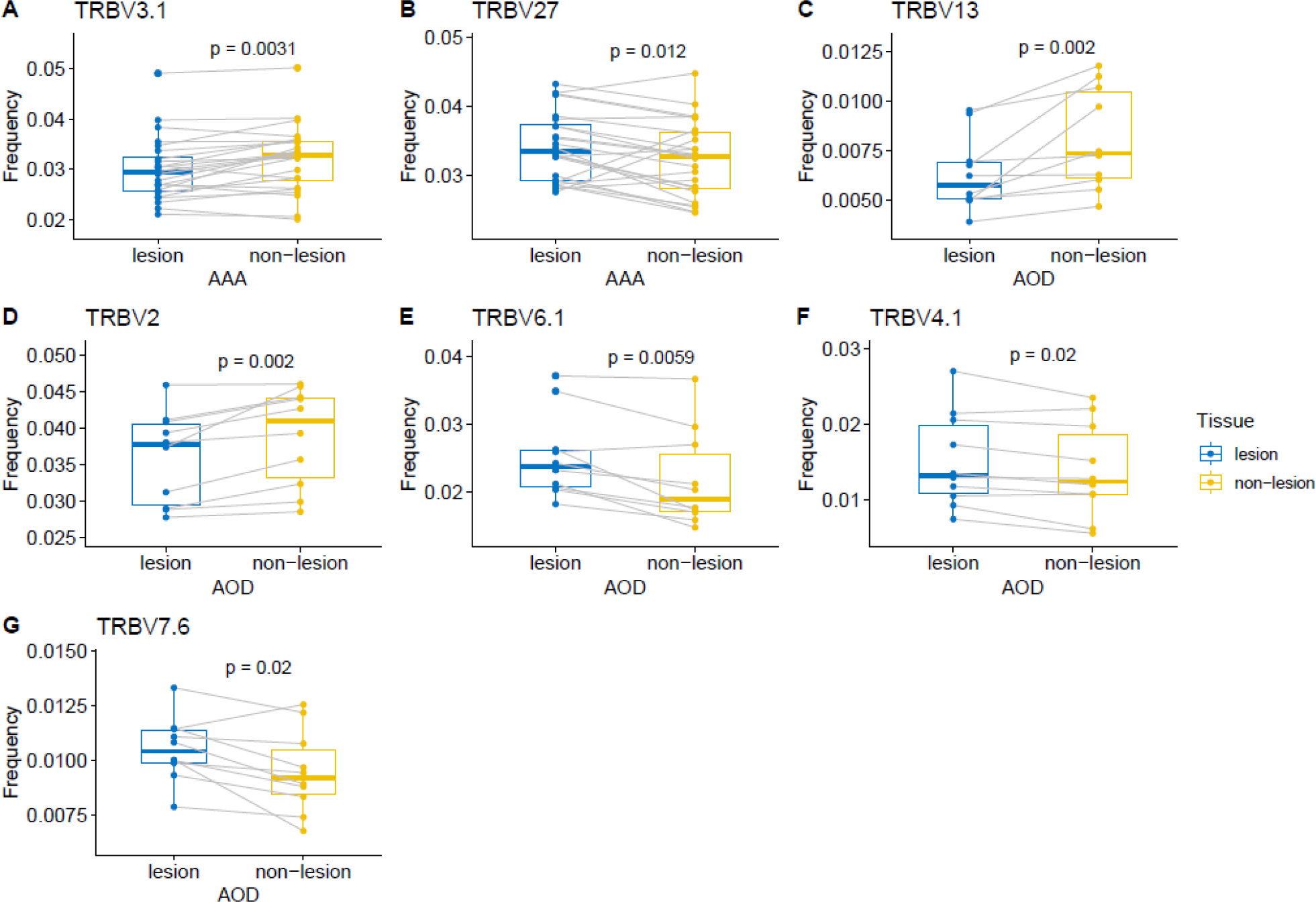
V segment usage differences in PVAT of lesion *vs*. non-lesion sites within AAA and AOD. Box plots show the different frequency of V gene segment usage for PVAT in lesion (blue) *vs*. non-lesion (yellow) for paired samples in AAA (**A** and **B**) and in AOD (**C-G**). A paired Wilcoxon test was used to assess the difference, with significance set at P<0.05.

## Discussion

This work is the first to investigate the clonal repertoire of T lymphocytes in PVAT of large vessels.

The overall results showed that an antigen-specific T-cell immune profile in PVAT characterizes and differentiates AAA from AOD, corroborating the idea of a different immune-pathogenic basis of the two vascular diseases.

Two main findings support our hypothesis. First, difference in TCRβ diversity estimate, together with the preferential use of the V/J gene segments, indicated the presence of a TCR bias as a consequence of a possible antigen-specific T-cell immune response in AAA compared to AOD. Second, we found disease-specific clonotypes that were uniquely present in a relevant fraction (~20%) of patients with AAA or AOD, and a small signature of 7 public clonotype that discriminated AAA patients from AOD patients with very high accuracy. As it is quite rare to observe identical clonotypes in multiple individuals, it is remarkable to find common clonotypes in such a large proportion of AAA or AOD patients and that only a few specific clonotypes can correctly distinguish patients with vascular diseases that share many risk factors but have their own epidemiological and pathogenic peculiarities.

Clonally expanded T-cells have already been reported in AAA lesions (Lu et al., 2014), although not yet at the PVAT level. However, the presence of non-shared clonally expanded T cells alone may not necessarily be related to the disease, but rather could be a likely consequence of an age-related condition, where clonal expansion is expected to increase in the elderly population (Britanova et al., 2014; Trofimov et al., 2022). This is consistent with the observed private hyperexpanded clonotypes in almost all AAA and AOD patients, who are typically of advanced age. However, we cannot rule out the possibility that the hyper-expansion of clonotypes in our patients was the result of an ongoing immune response to different AAA- and AOD-associated antigens.

The differences found between the TCRβ repertoire of small and large AAAs, in terms of differential V segment usage and public clonotype distribution, may suggest that the T-cell immune response evolves during disease progression, confirming what we had already hypothesized in our previous work (Piacentini et al., 2019). Although a putative autoantigen for AAA has been previously identified (Xia et al., 1996), we propose that during the course of the disease, as a result of tissue damage (e.g. extracellular matrix degradation) and cell death, a variety of (auto)antigens are generated and trigger a nuanced antigen-driven T-cell response.

Furthermore, we should also consider that the binding specificity of the TCR is not so strict due to TCR degeneracy, which implies that many TCRs can recognize the same antigen and many antigens can be recognized by the same TCR (Sewell, 2012). TCR degeneracy would thus broaden the spectrum of possible antigen-specific T-cell responses, which thus might present inter-individual peculiarities and consequently reduce the overlap of the overall TCRβ repertoire within the group of AAA patients. Taken together, these observations may explain why, in the context of AAA pathogenesis, the T-cell responses can adapt and vary as the disease progresses.

The results on V/J segment usage in AAA vs. AOD and within AAA subtypes (i.e. small vs. large) raised other intriguing, albeit speculative, considerations. The most significant differences in gene segment usage mainly involved the J segment genes in AAA vs. AOD and the V segment genes in small vs. large. The J segments determine part of the specificity of the CDR3 amino acid sequence and thus the epitope recognition. Differences in J segment frequency might therefore be associated with the ability to recognize a specific antigen/epitope and emerged as a more crucial characteristic when comparing AAA against AOD. The V segments, on the other hand, contribute to the amino acid sequence diversity of CDR3 and the binding to HLA molecules through the interaction with the CDR1 and CDR2, which are encoded by the V segment itself. The differences observed between small and large AAA only for the V segment genes suggest that the two subgroups of AAA patients may trigger a different T-cell immune response, which would depend on specific interactions of the TCR with certain HLA molecules expressed by APCs. This observation is supported by various studies showing significant associations between AAA and the expression of both specific class I and II HLA alleles, although these findings are controversial (Tilson et al., 1996; Tromp et al., 2006; Badger et al., 2007; Anaya-Ayala et al., 2019). There are several possible explanations for these inconclusive results: for instance, the immune response restricted to specific HLA molecules is not constant throughout the course of the disease, but may change and differentially characterize the early and late stages of disease progression, as we inferred by highlighting the observed differences between patients with small and large AAA.

The genetic component of susceptibility to AAA is still unclear. Although a certain percentage of patients have a family history of AAA (up to 15%), polymorphic variants or HLA alleles capable of discriminating those most at risk have not yet been identified (Pinard Amélie et al., 2019). In addition, the unclear associations between HLA alleles and AAA may argue against the autoimmune theory of AAA. It is therefore possible that only a certain number of individuals that combine a particular genetic background (e.g. HLA variants) and predisposing risk factors for an increased immune-inflammatory response in AAA (e.g. smoking, age) are actually susceptible to a process of chronic progression of the disease.

In this study, we demonstrated the specificity of the TCRβ repertoire in AAA and AOD by various means, including TCRβ diversity estimate, V/J segment usage, presence of common and disease-specific clonotypes, which taken together could be considered as an evidence of disease-related antigen-specific immune response. However, whether a pathogenic T-cell response or a lack of control of an autoimmune response (or a combination of the two) is responsible for the pathogenesis of AAA remains to be elucidated. In particular, a number of studies now suggest that a deficiency in regulatory T-cells function may increase inflammation and exacerbate AAA, and that this difference also occurs when compared to AOD (Yin et al., 2010; Li et al., 2022).

We must also point out that our work has some limitations. Studies on the diversity of the immune repertoire are complex, and only in recent years that next-generation sequencing technologies have made it possible to gain a deeper insight into the clonal repertoire of T cells. As a result, the analysis of immune repertoire sequencing data is also complex, and both the technology and the analytical methods to define and characterize the clonal repertoire have no gold standard, but are constantly evolving (Rosati et al., 2017; Heather et al., 2018; Bradley and Thomas, 2019). Identification of pathogenic T-cells in (auto)immune disease is further complicated by the fact that they represent a small percentage of all T-cells, even within the primary organ involved in the autoimmune process (Mitchell and Michels, 2020). The small number of patients constituting our cohort may limit the generalizability of the results obtained and, although our work is based on the collection of rare samples such as periaortic fat in patients with abdominal aortic pathology, our results would be strengthened by validation on a cohort of independent patients. Finally, we analyzed the PVAT transcriptome on whole tissue RNA extracts, which did not allow us to estimate the precise contribution of the different T-cell subtypes.

In conclusion, our results showed that an antigen-driven T-cell immune response characterized AAA and distinguished it from another type of aortic disease. This could occur as an initial response to environmental triggers that, in combination with risk factors, *e*.*g*. smoke (Tilson, 2017), in genetically susceptible individuals may lead to immunopathogenic T-cell responses that contribute to the development and the progression of the disease.

Our work provides new insights into the immunological component of PVAT in large-vessel diseases. We propose that T-cell immune response in PVAT is central in AAA pathogenesis and could be considered as a possible target for developing effective immunomodulant pharmacological treatment of the disease (AlZaim et al., 2020; Wang and Murphy, 2018; Serra and Santamaria, 2019). Furthermore, as it has been suggested that AAA may be a focal manifestation of a systemic disease (Langenskiöld et al., 2020), we can expect that a blood-based T-cell immunological profile could be used as a minimally invasive diagnostic tool to identify most at risk patients, just as in our study we have identified locally T-cell clonotypes that can discriminate with high accuracy patients with different aortic pathologies and those with different AAA progression.

## Methods

### Study population

This study included two groups of patients affected by abdominal aortic disease, including a group of 28 patients with an infrarenal AAA and a group of 11 patients with AOD, undergoing elective surgery at Centro Cardiologico Monzino between June 2010 and December 2014. **Table 2** summarizes the characteristics of AAA and AOD patients. For both patient groups, exclusion criteria from the study included Marfan syndrome, active or recent (5 years) cancer, recent major surgery (6 months), mycotic aneurysms, or disorders of the immune system, such as autoimmune diseases or vasculitis. Specifically, morphological features of the aneurysms (shape, thrombi content, position, length, and maximum diameter) were acquired from abdominal computed tomography scans performed within 2 months before surgery. The maximum diameter of the aneurysmal lesions ranged from 44.2 to 85.0 mm. Aneurysms with a maximum diameter ≥55.0 (n=7) or <55.0 mm (n=21) were defined as large and small, respectively. Elective repair of both large and small AAA was performed in compliance with the international and national guidelines for the treatment of AAA patients (Moll et al., 2011). Patients with AOD may present with varying degrees of obstruction at the site of the abdominal aorta. In accordance with the Trans-Atlantic Inter-Society Consensus Document on Management of Peripheral Arterial Disease (TASC) classification (Norgren et al., 2007), we described our AOD patients with type D lesions with either (i) infra-renal aortoiliac occlusions or (ii) unilateral or bilateral common iliac artery (CIA) occlusions or diffuse multiple stenoses involving the CIA with stenosis of the distal aorta. The description was made by detailed visual analysis of a contrast-enhanced tomography scan performed within two months prior to surgery. The study was approved by the Ethical Committee of Centro Cardiologico Monzino, and all the participants signed a written informed consent.

**Table 2.** Patient clinical characteristics. Categorical variables are shown as counts (n) and percentage (%); quantitative demographic variables are expressed both as mean ± standard deviation (SD) and as median and interquartile range [Q1, Q3]. Medications are summarized by drug class. Comparisons of AAA vs. AOD for categorical variables were made by Fisher’s exact test, when appropriate. Continuous variables were compared by two-tailed t-test or Wilcoxon test for non-normally distributed variables. Statistical significance for comparison: P<0.05. AAA indicates abdominal aortic aneurysm; ACE, angiotensin-converting-enzyme; AOD, Aortic Occlusive Disease; ARBs, angiotensin II receptor blockers; BMI, body mass index; CeVD, cerebrovascular disease; CHD, coronary heart disease; CKD, chronic kidney disease; HDL-c, high-density lipoprotein cholesterol.

|  | AAA<br>(N=28) | AOD<br>(N=11) | Overall<br>(N=39) | P-Val |
| --- | --- | --- | --- | --- |
| **Demographics** |  |  |  |  |
| **Sex** |  |  |  |  |
| M/F | 28/0 | 8/3 | 36/3 | **0.02** |
| **Age, y** |  |  |  |  |
| Mean ± SD | 68.0 ± 7.10 | 62.8 ± 8.00 | 66.6 ± 7.64 | 0.08 |
| Median [Q1, Q3] | 68.5 [62.0, 72.3] | 62.0 [58.0, 66.5] | 68.0 [59.5, 72.0] |  |
| **BMI, kg/m2** |  |  |  |  |
| Mean ± SD | 26.0 ± 2.59 | 26.3 ± 3.09 | 26.1 ± 2.71 | 0.81 |
| Median [Q1, Q3] | 25.8 [24.5, 27.3] | 25.5 [24.0, 28.2] | 25.8 [24.4, 27.5] |  |
| Missing | 2 (7.1%) | 0 (0%) | 2 (5.1%) |  |
| **CHD** | 12 (42.9%) | 5 (45.5%) | 17 (43.6%) | 0.70 |
| Missing | 0 (0%) | 2 (18.2%) | 2 (5.1%) |  |
| **CeVD** | 6 (21.4%) | 3 (27.3%) | 9 (23.1%) | 0.69 |
| **Hyperuricemia** | 1 (3.6%) | 1 (9.1%) | 2 (5.1%) | 0.50 |
| Missing | 1 (3.6%) | 0 (0%) | 1 (2.6%) |  |
| **T2DM** | 2 (7.1%) | 4 (36.4%) | 6 (15.4%) | 0.05 |
| Missing | 1 (3.6%) | 0 (0%) | 1 (2.6%) |  |
| **CKD** | 2 (7.1%) | 1 (9.1%) | 3 (7.7%) | 1 |
| Missing | 1 (3.6%) | 0 (0%) | 1 (2.6%) |  |
| **Risk factors** |  |  |  |  |
| **Smokers** | 8 (28.6%) | 5 (45.5%) | 13 (33.3%) | 0.46 |
| Missing | 1 (3.6%) | 0 (0%) | 1 (2.6%) |  |
| **Past Smokers** | 18 (64.3%) | 6 (54.5%) | 24 (61.5%) | 0.71 |
| Missing | 1 (3.6%) | 0 (0%) | 1 (2.6%) |  |
| **Hypertension** | 15 (53.6%) | 7 (63.6%) | 22 (56.4%) | 0.92 |
| Missing | 1 (3.6%) | 0 (0%) | 1 (2.6%) |  |
| **Dyslipidemia** | 17 (60.7%) | 8 (72.7%) | 25 (64.1%) | 0.84 |
| Missing | 1 (3.6%) | 0 (0%) | 1 (2.6%) |  |
| **Laboratory tests** |  |  |  |  |
| **Total cholesterol, mg/dL** |  |  |  |  |
| Mean ± SD | 194 ± 32.5 | 194 ± 53.9 | 194 ± 38.9 | 0.97 |
| Median [Q1, Q3] | 197 [171, 211] | 186 [168, 238] | 196 [170, 216] |  |
| **HDLc, mg/dL** |  |  |  |  |
| Mean ± SD | 40.9 ± 10.0 | 60.3 ± 38.8 | 46.4 ± 23.4 | 0.22 |
| Median [Q1, Q3] | 41.5 [35.3, 47.3] | 41.0 [40.0, 63.0] | 41.0 [36.5, 49.0] |  |
| **Triglycerides, mg/dL** |  |  |  |  |
| Mean ± SD | 132 ± 68.8 | 149 ± 63.3 | 137 ± 66.9 | 0.28 |
| Median [Q1, Q3] | 120 [84.8, 148] | 137 [116, 175] | 127 [88.5, 163] |  |
| **Medications** |  |  |  |  |
| **Antiaggregants** | 23 (82.1%) | 10 (90.9%) | 33 (84.6%) | 0.66 |
| **Diuretics** | 5 (17.9%) | 4 (36.4%) | 9 (23.1%) | 0.24 |
| **ACE inhibitors** | 5 (17.9%) | 1 (9.1%) | 6 (15.4%) | 0.66 |
| **Statins** | 11 (39.3%) | 5 (45.5%) | 16 (41.0%) | 0.73 |
| **Beta blockers** | 9 (32.1%) | 3 (27.3%) | 12 (30.8%) | 1 |
| **Calcium blockers** | 7 (25.0%) | 2 (18.2%) | 9 (23.1%) | 1 |
| **Proton pump inhibitors** | 12 (42.9%) | 6 (54.5%) | 18 (46.2%) | 0.72 |
| **Oral hypoglycemics** | 2 (7.1%) | 2 (18.2%) | 4 (10.3%) | 0.56 |
| **ARBs** | 5 (17.9%) | 4 (36.4%) | 9 (23.1%) | 0.24 |

### Sample Collection and RNA Isolation

At the time of surgery, samples of periaortic fat (PVAT) surrounding the lesion site of the abdominal aorta (aneurysm or atherosclerotic plaques or thrombus) and periaortic fat obtained from the aortic neck (non-lesioned segment), proximal to the lesion site, were obtained. Care was taken to avoid areas with visible fibrous tissue or vessels. PVAT samples were immediately snap frozen in liquid nitrogen and stored at -80°C until processing. Total RNA was extracted from 50 to 100 mg of frozen PVAT samples using TRIzol reagent (Life Technologies) and treated with TURBO DNase (Thermo Fisher Scientific) to remove genomic contamination, according to the manufacturer’s instructions. RNA isolation replicates from the PVAT surrounding the lesion site of the abdominal aorta of two subjects (namely, #01 and #03) were obtained by a second round of RNA extraction in a different time period following the same RNA isolation protocol. The Infinite M200 PRO multimode microplate reader (Tecan) and the 2100 Bioanalyzer (Agilent Technologies) were used to assess RNA yield/purity and RNA integrity, respectively. Total RNA was available from 28 dilated and 24 non-dilated segments of the abdominal aorta from AAA and from 11 atherosclerotic and 10 non-atherosclerotic segments of the abdominal aorta from AOD.

### T-cell receptor β-chain (TCRβ) sequencing

Two-hundred and fifty ng of total RNA was reverse-transcribed to cDNA using the SuperScript™ IV VILO™ Kit (Thermo Fisher Scientific). Replicates of PVAT surrounding the lesion site in the abdominal aorta of two subjects (namely, #36 and #41) were also reverse transcribed starting from 100 and 500 ng of total RNA. cDNA was amplified using the Oncomine TCR Beta-LR Assay (Thermo Fisher Scientific) and barcoded library were prepared using Ion Ampliseq library kit Plus (Thermo Fisher Scientific) and Ion Select Barcodes, according to manufacturer’s instructions. The Oncomine TCR Beta-LR Assay uses multiplexed primers targeting the F1 region of the variable V segment gene and the constant segment gene (64 and 2 primers respectively) of the TCRβ-VDJ rearrangement in cDNA, producing an amplicon of ~330 base pairs and allowing sequencing of all three complementarity determining regions (CDR1, CDR2 and CDR3). Libraries were then purified using Agencourt AMPure XP beads (Beckman Coulter), eluted in 50 µL and quantified through the Ion Quantitation kit (Thermo Fisher Scientific). Once the libraries were diluted to 25 pM with Low TE Buffer, 6 to 8 samples were pooled and loaded on the Ion 530™ Chip (Thermo Fisher Scientific) using the Ion Chef™ Instrument (Thermo Fisher Scientific). Sequencing was than performed by the Ion GeneStudio™ S5 Prime System (Thermo Fisher Scientific).

### Bioinformatic analysis

#### Data preparation

MiXCR v3.0.12 software (Bolotin et al., 2015) was used to align and assemble clonotypes from bulk raw sequencing data (fastq file format) as produced by Ion GeneStudio™ S5 Prime System. MiXCR parameters are specified in the **Supplementary Material 1**. The ‘clonotype tables’ generated for each sample were used for subsequent analysis through the VDJtools v1.2.1 software (Shugay et al., 2015), which required appropriate clonotype table conversion by the *Convert* function of VDJtools. TCRβ sequencing can present technical issue that may affect downstream analysis and results. Cross-sample contamination can indeed occur at library preparation or sequencing stage and lead to a number of artificial shared clonotypes among samples sequenced in the same batch. Thus, putative batch effects were evaluated on raw data through all-versus-all pairwise overlap of samples and hierarchical clustering on computed repertoire similarity measures. Samples presented a clear chip sequencing batch effect so that a normalization pre-processing, which included a ‘decontamination’ and a ‘down-sampling’ procedure for limiting the effect of both PCR contamination and biases for sequencing depth, was applied. For this purpose, the *Decontaminate* and *DownSample* functions of VDJtools were used (see parameter details in the **Supplementary Material 1**). To ensure a wiser trade-off between data normalization and possible overcorrection, the ‘decontamination’ step was performed, with different values of the ‘Parent-to-child clonotype frequency ratio for contamination filtering’ (option -r for *Decontaminate*), separately on samples within each sequencing chip batch, to prevent the exclusion of true positive clonotypes from samples sequenced on different chips. The threshold of the number of reads/clonotypes to be taken for ‘down-sampling’ was instead chosen by evaluating the rarefaction curves on the raw data of all samples (**Supplementary Figure S7**). Finally, the combined effect of tuning of the decontamination and down-sampling parameters was again evaluated by inspecting the hierarchical clustering on the pairwise overlap measures of samples until it was visualized that the observed sequencing batch effects had been substantially removed. The final thresholds were set at 2000 for the ‘Parent-to-child clonotype frequency ratio for contamination filtering’ and 250000 ‘reads/clonotypes’ for the *Decontaminate* and *DownSample* functions, respectively.

#### Data analysis

Normalized clonotype tables were used as input for data exploration and downstream statistical analysis. Diversity estimation was assessed through the *CalcDiversityStats* function of VDJtools, which include several diversity statistics. Notably, a lower bound estimate of total repertoire diversity (*i*.*e*. the Chao-1 estimate) was preferentially used because lower bound estimates were successfully applied in studies for the comparison of TCR repertoire diversity under uneven sample sizes, as it is our case (Shugay et al., 2015; Robins et al., 2009; Qi et al., 2014).

Repertoire overlap analysis was performed by running two VDJtools functions: *CalcPairwiseDistances* for calculating pairwise overlap between samples, and *ClusterSamples* for hierarchical clustering of those measures. The clonotype-wise sum of geometric mean frequencies calculated as:

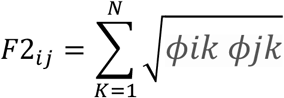

was implemented as the similarity measure for clonotypes that matched identical CDR3 amino acid sequence and V segment (*i*.*e*. aaV). F2 provides slightly more robust results compared to other similarity measures in case cross-sample contamination still occurs (Shugay et al., 2015).

The frequency of associated reads for each of V/J segments (aka segment usage) of each sample was calculated through the *CalcSegmentUsage* of VDJtools.

Homeostatic space was estimated through the *repClonality* function of the immunarch R package (Nazarov et al., 2022), identifying clonotypes as Rare ≤0.00001; 0.00001<Small≤0.0001; 0.0001<Medium≤0.001; 0.001<Large≤0.01; Hyperexpanded > 0.01 (default boundaries).

A repertoire of per-sample proportions of ‘public’ clonotypes identified as coding-only sequences using CDR3 amino acid sequences and V genes was generated through the *pubRep* function of immunarch software. A clonotype was considered ‘public’ if that clonotype was shared by at least 2 samples regardless of disease phenotype, *i*.*e*. the clonotype could be shared both within AAA or AOD or between any patients of the two disease groups. On the contrary, clonotypes that are not shared with any sample are defined as ‘private’.

The public clonotypes (defined by aaV) were then used to identify disease-associated clonotypes by first constructing a contingency table of clonotype(s) (present/absent) per disease group (AAA/AOD) and then by comparing the proportions of clonotypes between AAA and AOD using Fisher’s exact test. A clonotype was regarded as disease-specific if the Fisher’s exact test was significant at P-Value<0.05.

The resulting set of disease-associated clonotypes was assessed for its capacity to discriminate between AAA from AOD patients of our study population. A recursive feature elimination was first applied to identify a smaller set of disease-associated clonotypes for AAA and AOD sample discrimination and then different classification models were developed to test both linear and non-linear methods (caret R package). To obtain more robust results, up-sampling and 10-times repeated, 5-fold cross validation were also applied to adjust for class imbalance and to set appropriate hyperparameters, respectively. The area under the receiver operating characteristic curve (AUC-ROC) was used to select the optimal model with the largest value and to evaluate the overall classification performance along with sensitivity and sensibility.

Support vector machine with radial basis function kernel (svmRadial method in caret R package), a non-linear method, outperformed the other models displaying a higher accuracy, sensitivity, and specificity for disease discrimination and was chosen as the optimal classifier.

Details of the parameters of the methods used in the bioinformatic workflow are reported in the online-only **Supplementary Material 1**.

### Statistics

All the statistical analyses were performed in the R environment v4.1.1. Demographic and clinical categorical data are presented as counts and percent, continuous data as the median and interquartile range (Q1–Q3). Categorical and continuous variables were compared between AAA *vs*. AOD or large *vs*. small AAA subgroups by chi-squared (χ2) and Wilcoxon rank sum (two-sided) test, respectively.

Wilcoxon rank sum test with continuity correction was also performed to test differences of diversity measures, V/J segment usage and homeostatic space (rare to hyper-expanded proportions) variables between AAA and AOD and between small and large AAA. A P-Value < 0.05 was considered statistically significant.

For paired-samples analysis between PVAT of the lesion and non-lesion segment of the aorta a paired Wilcoxon test was used for TCRβ diversity estimate, V/J segment usage and homeostatic space variables, and the McNemar’s test for the pairwise testing of public clonotype proportions. A P-Value<0.05 was considered as significant.

## Supplementary information

## Funding

This work was supported by the Italian Ministry of Health (Research Projects RC Nos. 2600658, 2627621, and 2631196).

## Financial conflicts of interest

The authors declare no conflict of interests.

## Abbreviations

AAA: abdominal aortic aneurysm
AOD: aortic occlusive disease
AUC: area under the (ROC) curve
APC: antigen presenting cells
CDR: complementarity determining region
HLA: human leukocyte antigen
PVAT: perivascular adipose tissue
ROC: receiver operating characteristic
TCRβ: T-cell receptor β-chain

## Notes

### Competing Interest Statement

The authors have declared no competing interest.

## References

AlZaim, I., S.H. Hammoud, H. Al-Koussa, A. Ghazi, A.H. Eid, and A.F. El-Yazbi. 2020. Adipose Tissue Immunomodulation: A Novel Therapeutic Approach in Cardiovascular and Metabolic Diseases. Front. Cardiovasc. Med. 7:602088. doi:10.3389/fcvm.2020.602088.

Anaya-Ayala, J.E., S. Hernandez-Doño, M. Escamilla-Tilch, J. Marquez-Garcia, K. Hernandez-Sotelo, R. Lozano-Corona, D. Ruiz-Gomez, J. Granados, and C.A. Hinojosa. 2019. Genetic polymorphism of HLA-DRB1 alleles in Mexican mestizo patients with abdominal aortic aneurysms. BMC Med. Genet. 20:102. doi:10.1186/s12881-019-0833-8.

Badger, S.A., C.V. Soong, M.E. O’Donnell, and D. Middleton. 2007. The role of human leukocyte antigen genes in the formation of abdominal aortic aneurysms. J. Vasc. Surg. 45:475–480. doi:10.1016/j.jvs.2006.09.067.

Biros, E., G. Gäbel, C.S. Moran, C. Schreurs, J.H.N. Lindeman, P.J. Walker, M. Nataatmadja, M. West, L.M. Holdt, I. Hinterseher, C. Pilarsky, and J. Golledge. 2015. Differential gene expression in human abdominal aortic aneurysm and aortic occlusive disease. Oncotarget. 6:12984–12996. doi:10.18632/oncotarget.3848.

Bolotin, D.A., S. Poslavsky, I. Mitrophanov, M. Shugay, I.Z. Mamedov, E.V. Putintseva, and D.M. Chudakov. 2015. MiXCR: software for comprehensive adaptive immunity profiling. Nat. Methods. 12:380–381. doi:10.1038/nmeth.3364.

Bradley, P., and P.G. Thomas. 2019. Using T Cell Receptor Repertoires to Understand the Principles of Adaptive Immune Recognition. Annu. Rev. Immunol. 37:547–570. doi:10.1146/annurev-immunol-042718-041757.

Chowdhury, R.R., J. D’Addabbo, X. Huang, S. Veizades, K. Sasagawa, D.M. Louis, P. Cheng, J. Sokol, A. Jensen, A. Tso, V. Shankar, B.S. Wendel, I. Bakerman, G. Liang, T. Koyano, R. Fong, A.N. Nau, H. Ahmad, J. Gopakumar, R. Wirka, A.S. Lee, J. Boyd, Y.J. Woo, T. Quertermous, G.S. Gulati, S. Jaiswal, Y.-H. Chien, C.K.F. Chan, M.M. Davis, and P.K. Nguyen. 2022. Human Coronary Plaque T Cells Are Clonal and Cross-React to Virus and Self. Circ. Res. 130:1510–1530. doi:10.1161/CIRCRESAHA.121.320090.

Criqui, M.H., and V. Aboyans. 2015. Epidemiology of peripheral artery disease. Circ. Res. 116:1509–1526. doi:10.1161/CIRCRESAHA.116.303849.

Farmanbar, A., R. Kneller, and S. Firouzi. 2019. RNA sequencing identifies clonal structure of T-cell repertoires in patients with adult T-cell leukemia/lymphoma. Npj Genomic Med. 4:1–9. doi:10.1038/s41525-019-0084-9.

Folkesson, M., E. Vorkapic, E. Gulbins, L. Japtok, B. Kleuser, M. Welander, T. Länne, and D. Wågsäter. 2017. Inflammatory cells, ceramides, and expression of proteases in perivascular adipose tissue adjacent to human abdominal aortic aneurysms. J. Vasc. Surg. 65:1171–1179.e1. doi:10.1016/j.jvs.2015.12.056.

Golledge, J. 2019. Abdominal aortic aneurysm: update on pathogenesis and medical treatments. Nat. Rev. Cardiol. 16:225–242. doi:10.1038/s41569-018-0114-9.

Guzik, T.J., D.S. Skiba, R.M. Touyz, and D.G. Harrison. 2017. The role of infiltrating immune cells in dysfunctional adipose tissue. Cardiovasc. Res. 113:1009–1023. doi:10.1093/cvr/cvx108.

Heather, J.M., M. Ismail, T. Oakes, and B. Chain. 2018. High-throughput sequencing of the T-cell receptor repertoire: pitfalls and opportunities. Brief. Bioinform. 19:554–565. doi:10.1093/bib/bbw138.

Horimatsu, T., H.W. Kim, and N.L. Weintraub. 2017. The Role of Perivascular Adipose Tissue in Non-atherosclerotic Vascular Disease. Front. Physiol. 8:969. doi:10.3389/fphys.2017.00969.

Huang Cao, Z.F., E. Stoffel, and P. Cohen. 2017. Role of Perivascular Adipose Tissue in Vascular Physiology and Pathology. Hypertens. Dallas Tex 1979. 69:770–777. doi:10.1161/HYPERTENSIONAHA.116.08451.

Kent, K.C., R.M. Zwolak, N.N. Egorova, T.S. Riles, A. Manganaro, A.J. Moskowitz, A.C. Gelijns, and G. Greco. 2010. Analysis of risk factors for abdominal aortic aneurysm in a cohort of more than 3 million individuals. J. Vasc. Surg. 52:539–548. doi:10.1016/j.jvs.2010.05.090.

Langenskiöld, M., K. Smidfelt, J. Nordanstig, G. Bergström, and Å. Tivesten. 2020. Leukocyte subsets and abdominal aortic aneurysms detected by screening in men. J. Intern. Med. 288:345–355. doi:10.1111/joim.13040.

Li, J., N. Xia, D. Li, S. Wen, S. Qian, Y. Lu, M. Gu, T. Tang, J. Jiao, B. Lv, S. Nie, D. Hu, Y. Liao, X. Yang, G. Shi, and X. Cheng. 2022. Aorta Regulatory T Cells with a Tissue-Specific Phenotype and Function Promote Tissue Repair through Tff1 in Abdominal Aortic Aneurysms. Adv. Sci. 9:2104338. doi:10.1002/advs.202104338.

Lu, S., J.V. White, W.L. Lin, X. Zhang, C. Solomides, K. Evans, N. Ntaoula, I. Nwaneshiudu, J. Gaughan, D.S. Monos, E.L. Oleszak, and C.D. Platsoucas. 2014. Aneurysmal Lesions of Patients with Abdominal Aortic Aneurysm Contain Clonally Expanded T Cells. J. Immunol. 192:4897–4912. doi:10.4049/jimmunol.1301009.

Mitchell, A.M., and A.W. Michels. 2020. T cell receptor sequencing in autoimmunity. J. Life Sci. Westlake Village Calif. 2:38–58. doi:10.36069/jols/20201203.

Moll, F.L., J.T. Powell, G. Fraedrich, F. Verzini, S. Haulon, M. Waltham, J.A. van Herwaarden, P.J.E. Holt, J.W. van Keulen, B. Rantner, F.J.V. Schlösser, F. Setacci, J.-B. Ricco, and European Society for Vascular Surgery. 2011. Management of abdominal aortic aneurysms clinical practice guidelines of the European society for vascular surgery. Eur. J. Vasc. Endovasc. Surg. Off. J. Eur. Soc. Vasc. Surg. 41 Suppl 1:S1–S58. doi:10.1016/j.ejvs.2010.09.011.

Nazarov, V.I., V.O. Tsvetkov, E. Rumynskiy, A.A. Popov, I. Balashov, and M. Samokhina. 2022. immunarch: Bioinformatics Analysis of T-Cell and B-Cell Immune Repertoires.

Norgren, L., W.R. Hiatt, J.A. Dormandy, M.R. Nehler, K.A. Harris, and F.G.R. Fowkes. 2007. Inter-Society Consensus for the Management of Peripheral Arterial Disease (TASC II). J. Vasc. Surg. 45:S5–S67. doi:10.1016/j.jvs.2006.12.037.

Piacentini, L., M. Chiesa, and G.I. Colombo. 2020a. Gene Regulatory Network Analysis of Perivascular Adipose Tissue of Abdominal Aortic Aneurysm Identifies Master Regulators of Key Pathogenetic Pathways. Biomedicines. 8. doi:10.3390/biomedicines8080288.

Piacentini, L., C. Saccu, E. Bono, E. Tremoli, R. Spirito, G.I. Colombo, and J.P. Werba. 2020b. Gene-expression profiles of abdominal perivascular adipose tissue distinguish aortic occlusive from stenotic atherosclerotic lesions and denote different pathogenetic pathways. Sci. Rep. 10:6245. doi:10.1038/s41598-020-63361-5.

Piacentini, L., C. Vavassori, and G.I. Colombo. 2022. Trained Immunity in Perivascular Adipose Tissue of Abdominal Aortic Aneurysm—A Novel Concept for a Still Elusive Disease. Front. Cell Dev. Biol. 10.

Piacentini, L., J.P. Werba, E. Bono, C. Saccu, E. Tremoli, R. Spirito, and G.I. Colombo. 2019. Genome-Wide Expression Profiling Unveils Autoimmune Response Signatures in the Perivascular Adipose Tissue of Abdominal Aortic Aneurysm. Arterioscler. Thromb. Vasc. Biol. 39:237–249. doi:10.1161/ATVBAHA.118.311803.

Pinard Amélie, Jones Gregory T., and Milewicz Dianna M. 2019. Genetics of Thoracic and Abdominal Aortic Diseases. Circ. Res. 124:588–606. doi:10.1161/CIRCRESAHA.118.312436.

Qi, Q., Y. Liu, Y. Cheng, J. Glanville, D. Zhang, J.-Y. Lee, R.A. Olshen, C.M. Weyand, S.D. Boyd, and J.J. Goronzy. 2014. Diversity and clonal selection in the human T-cell repertoire. Proc. Natl. Acad. Sci. 111:13139–13144. doi:10.1073/pnas.1409155111.

Queiroz, M., and C.M. Sena. 2020. Perivascular adipose tissue in age-related vascular disease. Ageing Res. Rev. 59:101040. doi:10.1016/j.arr.2020.101040.

Robins, H.S., P.V. Campregher, S.K. Srivastava, A. Wacher, C.J. Turtle, O. Kahsai, S.R. Riddell, E.H. Warren, and C.S. Carlson. 2009. Comprehensive assessment of T-cell receptor β-chain diversity in αβ T cells. Blood. 114:4099–4107. doi:10.1182/blood-2009-04-217604.

Rosati, E., C.M. Dowds, E. Liaskou, E.K.K. Henriksen, T.H. Karlsen, and A. Franke. 2017. Overview of methodologies for T-cell receptor repertoire analysis. BMC Biotechnol. 17:61. doi:10.1186/s12896-017-0379-9.

Sagan, A., T.P. Mikolajczyk, W. Mrowiecki, N. MacRitchie, K. Daly, A. Meldrum, S. Migliarino, C. Delles, K. Urbanski, G. Filip, B. Kapelak, P. Maffia, R. Touyz, and T.J. Guzik. 2019. T Cells Are Dominant Population in Human Abdominal Aortic Aneurysms and Their Infiltration in the Perivascular Tissue Correlates With Disease Severity. Front. Immunol. 10:1979. doi:10.3389/fimmu.2019.01979.

Serra, P., and P. Santamaria. 2019. Antigen-specific therapeutic approaches for autoimmunity. Nat. Biotechnol. 37:238–251. doi:10.1038/s41587-019-0015-4.

Sewell, A.K. 2012. Why must T cells be cross-reactive? Nat. Rev. Immunol. 12:669–677. doi:10.1038/nri3279.

Shugay, M., D.V. Bagaev, M.A. Turchaninova, D.A. Bolotin, O.V. Britanova, E.V. Putintseva, M.V. Pogorelyy, V.I. Nazarov, I.V. Zvyagin, V.I. Kirgizova, K.I. Kirgizov, E.V. Skorobogatova, and D.M. Chudakov. 2015. VDJtools: Unifying Post-analysis of T Cell Receptor Repertoires. PLOS Comput. Biol. 11:e1004503. doi:10.1371/journal.pcbi.1004503.

Suh, M.K., R. Batra, J.S. Carson, W. Xiong, M.A. Dale, T. Meisinger, C. Killen, J. Mitchell, and B.T. Baxter. 2020. Ex vivo expansion of regulatory T cells from abdominal aortic aneurysm patients inhibits aneurysm in humanized murine model. J. Vasc. Surg. 72:1087–1096.e1. doi:10.1016/j.jvs.2019.08.285.

Tang, T.-T., Y.-C. Zhu, N.-G. Dong, S. Zhang, J. Cai, L.-X. Zhang, Y. Han, N. Xia, S.-F. Nie, M. Zhang, B.-J. Lv, J. Jiao, X.-P. Yang, Y. Hu, Y.-H. Liao, and X. Cheng. 2019. Pathologic T-cell response in ischaemic failing hearts elucidated by T-cell receptor sequencing and phenotypic characterization. Eur. Heart J. 40:3924–3933. doi:10.1093/eurheartj/ehz516.

Tilson, M.D. 2017. Autoimmunity in the Abdominal Aortic Aneurysm and its Association with Smoking. Aorta Stamford Conn. 5:159–167. doi:10.12945/j.aorta.2017.17.693.

Tilson, M.D., K.J. Ozsvath, H. Hirose, and S. Xia. 1996. A Genetic Basis for Autoimmune Manifestations in the Abdominal Aortic Aneurysm Resides in the MHC Class II Locus DR-beta-1a. Ann. N. Y. Acad. Sci. 800:208–215. doi:10.1111/j.1749-6632.1996.tb33311.x.

Tromp, G., T. Ogata, L. Gregoire, K.A.B. Goddard, M. Skunca, W.D. Lancaster, A.R. Parrado, Q. Lu, H. Shibamura, N. Sakalihasan, R. Limet, G.L. Mackean, C. Arthur, T. Sueda, and H. Kuivaniemi. 2006. HLA-DQA is associated with abdominal aortic aneurysms in the Belgian population. Ann. N. Y. Acad. Sci. 1085:392–395. doi:10.1196/annals.1383.045.

Wang, S.K., and M.P. Murphy. 2018. Immune Modulation as a Treatment for Abdominal Aortic Aneurysms. Circ. Res. 122:925–927. doi:10.1161/CIRCRESAHA.118.312870.

Xia, S., K. Ozsvath, H. Hirose, and M.D. Tilson. 1996. Partial amino acid sequence of a novel 40-kDa human aortic protein, with vitronectin-like, fibrinogen-like, and calcium binding domains: aortic aneurysm-associated protein-40 (AAAP-40) [human MAGP-3, proposed]. Biochem. Biophys. Res. Commun. 219:36–39. doi:10.1006/bbrc.1996.0177.

Yin, M., J. Zhang, Y. Wang, S. Wang, D. Böckler, Z. Duan, and S. Xin. 2010. Deficient CD4+CD25+ T regulatory cell function in patients with abdominal aortic aneurysms. Arterioscler. Thromb. Vasc. Biol. 30:1825–1831. doi:10.1161/ATVBAHA.109.200303.

Zhou, Y., W. Wu, J.S. Lindholt, G.K. Sukhova, P. Libby, X. Yu, and G.-P. Shi. 2015. Regulatory T cells in human and angiotensin II-induced mouse abdominal aortic aneurysms. Cardiovasc. Res. 107:98–107. doi:10.1093/cvr/cvv119.

