## Supplementary Material for "Deciphering abdominal aortic diseases through T-cell clonal repertoire of perivascular adipose tissue"

#### Details on parameters of software used in the bioinformatic workflow.

Herein, we report specific parameters used for the bioinformatic analysis.

MiXCR v3.0.12 parameters for alignment and clonotype assembly:

```
java -Xmx12g -Xms10g -jar mixcr.jar \
analyze amplicon \
-s hsa \
--starting-material rna \
--5-end v-primers \
--3-end c-primers \
--adapters adapters-present \
--assemble "-OseparateByV=true" \
--assemble "-OseparateByJ=true" \
--assemble "-OseparateByC=true" \
--contig-assembly \
--receptor-type trb \
```

VDJtools v1.2.1 parameters:

```
# conversion from MiXCR format:
java -Xmx16G -jar vdjtools-1.2.1.jar \
Convert \
-S mixcr
```

```
# decontamination:
java -Xmx16G -jar vdjtools-1.2.1.jar \
Decontaminate \
 --read-based \
-r 2000 \

# down-sampling:
java -Xmx16G -jar vdjtools-1.2.1.jar \
DownSample \
-x 250000 \
```

Options used for other VDJtools functions are: *CalcDiversityStats*: -i aaV; *CalcPairwiseDistances*: -i aaV and -p; *ClusterSamples*: -e F2 and -i aaV; *CalcSegmentUsage*: -u

### Supplementary Figures.

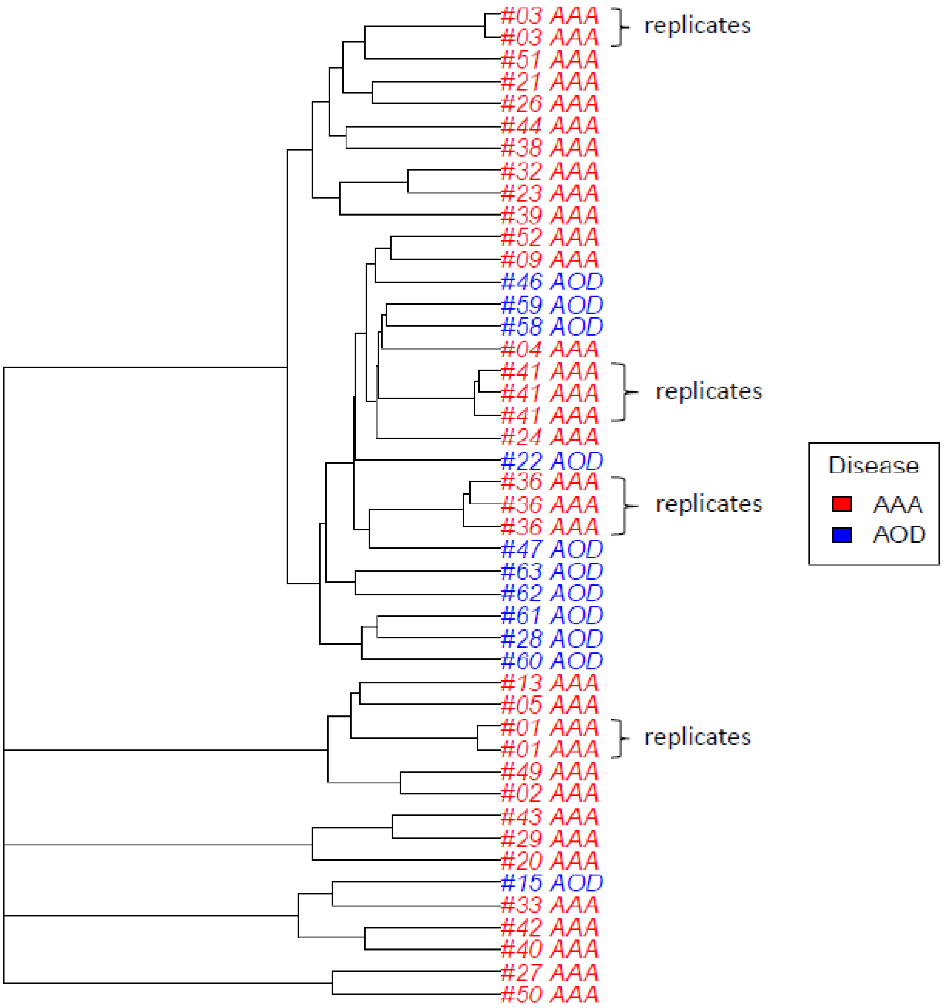


**Supplementary Figure S1. TCRβ repertoire overlapping in PVAT surrounding aortic lesion in AAA and AOD samples.** Hierarchical clustering of the clonotype similarity measure, calculated as the clonotype sum of geometric mean frequencies (F2; see Methods section of the manuscript) including sample replicates, is shown. Clonotypes were defined as those with identical CDR3 amino acid sequence and V segment. The three replicates of samples #41 and #36 were obtained using 100, 250 and 500 ng of total RNA input for sequencing, whereas the replicates of samples #01 and #03 were obtained from two different RNA extraction batches performed in two different time periods. The two sample #01 replicates were sequenced by using 100 and 250 ng of total RNA, whereas the two sample #03 replicates were sequenced using the same amount (250 ng) of total RNA. The clustering shows that the sample replicates are closely grouped, giving confidence in the correctness of the choice of metrics and the result obtained.

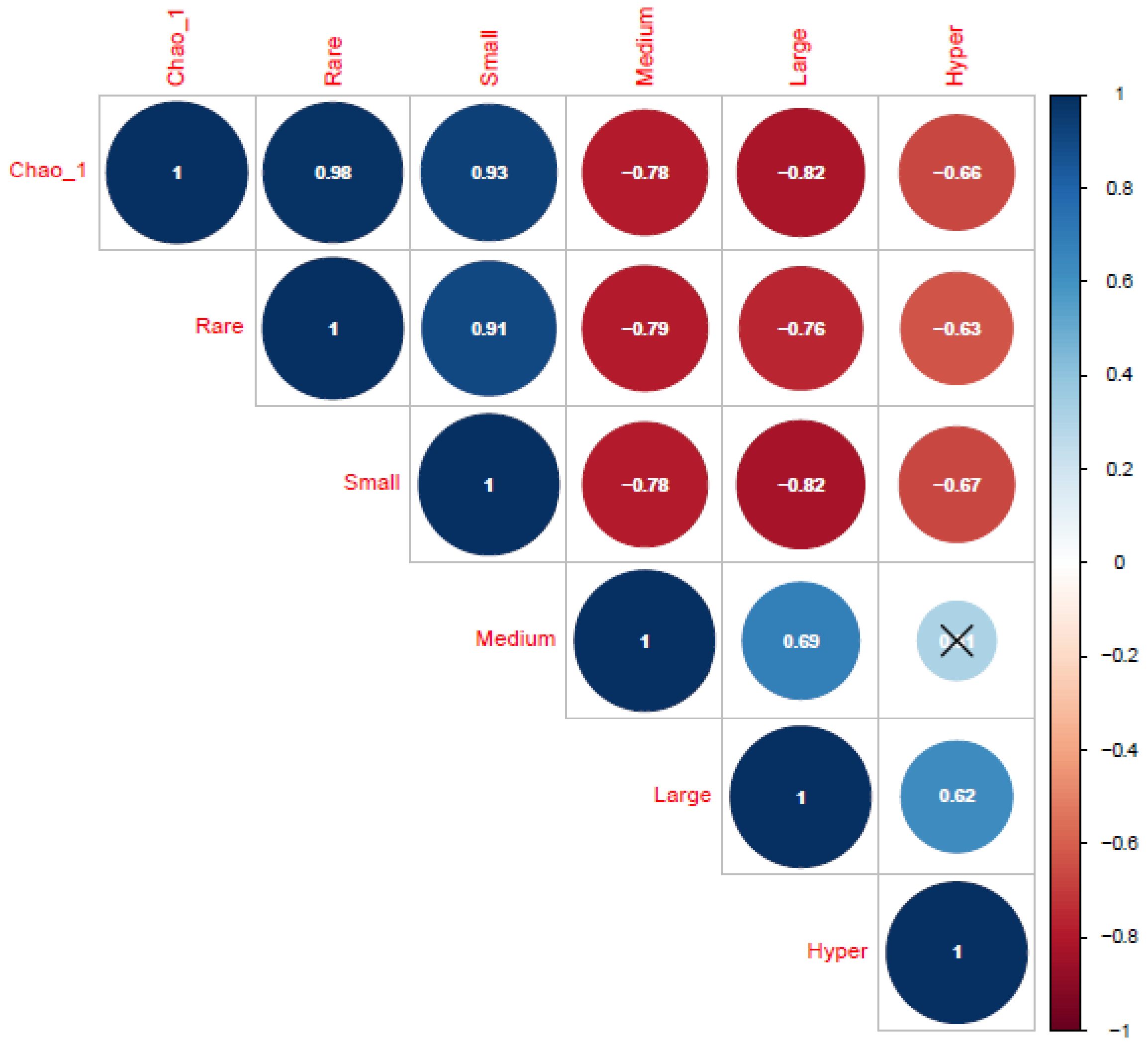


**Supplementary Figure S2. Correlation plot between diversity estimate and clonotype frequencies.** The graph shows Spearman's correlation between the Chao-1 diversity estimate and the relative frequency of clonotypes defined as Rare ≤0.00001; 0.00001<Small≤0.0001; 0.0001<Medium≤0.001; 0.001<Large≤0.01; and Hyperexpanded > 0.01. The degree of correlation coefficient (rho) is highlighted by a colour gradient from 1 (dark blue) to -1 (dark red). Significance was set at P-value<0.05. Non-significant correlations are marked by black crosses.

A

B

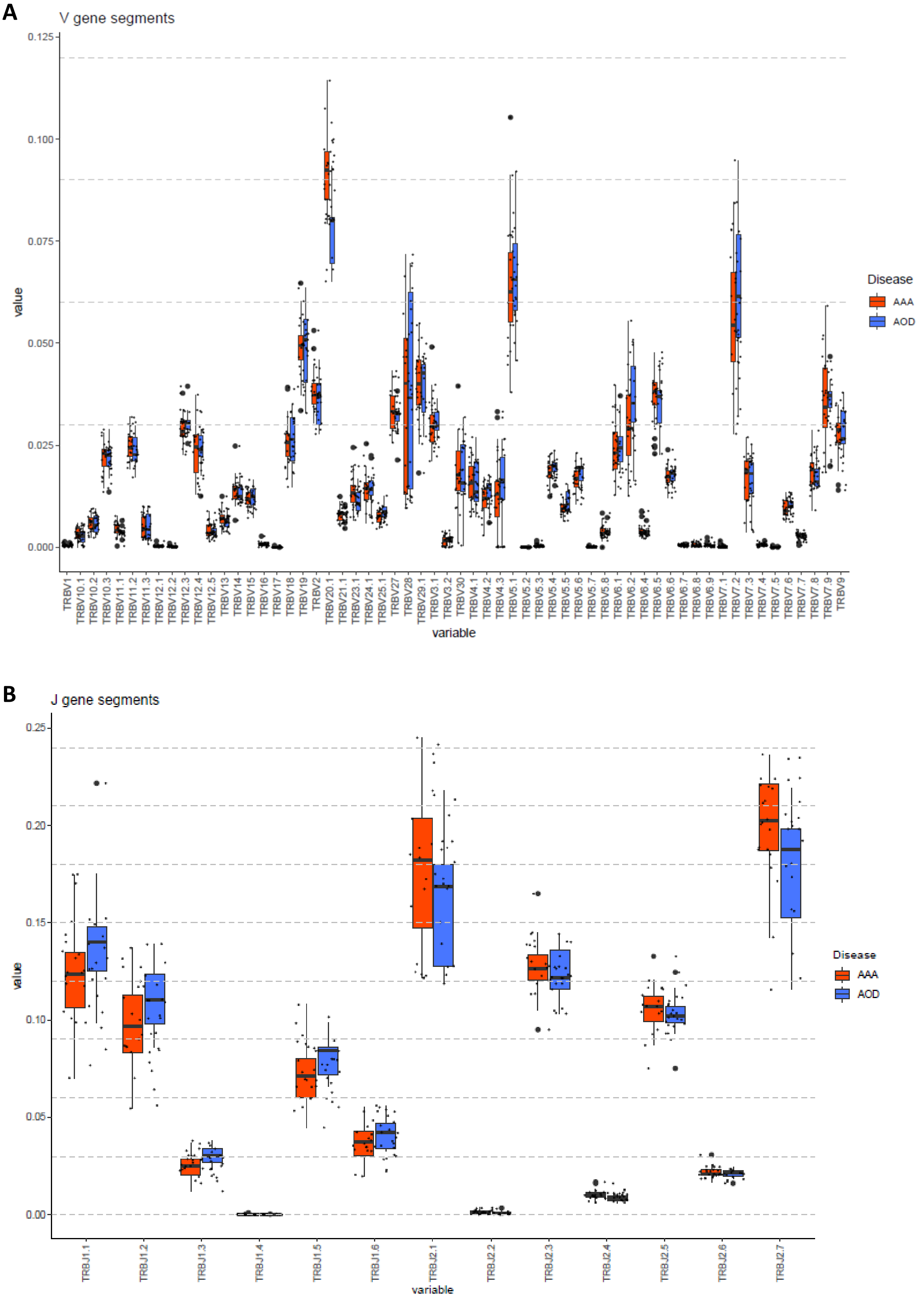


**Supplementary Figure S3. Overall V/J segment usage differences between AAA and AOD.** Box plots of the frequency of use of all V (**A**) and J (**B**) segments tested in the AAA *vs.* AOD comparison are shown in the same graph.

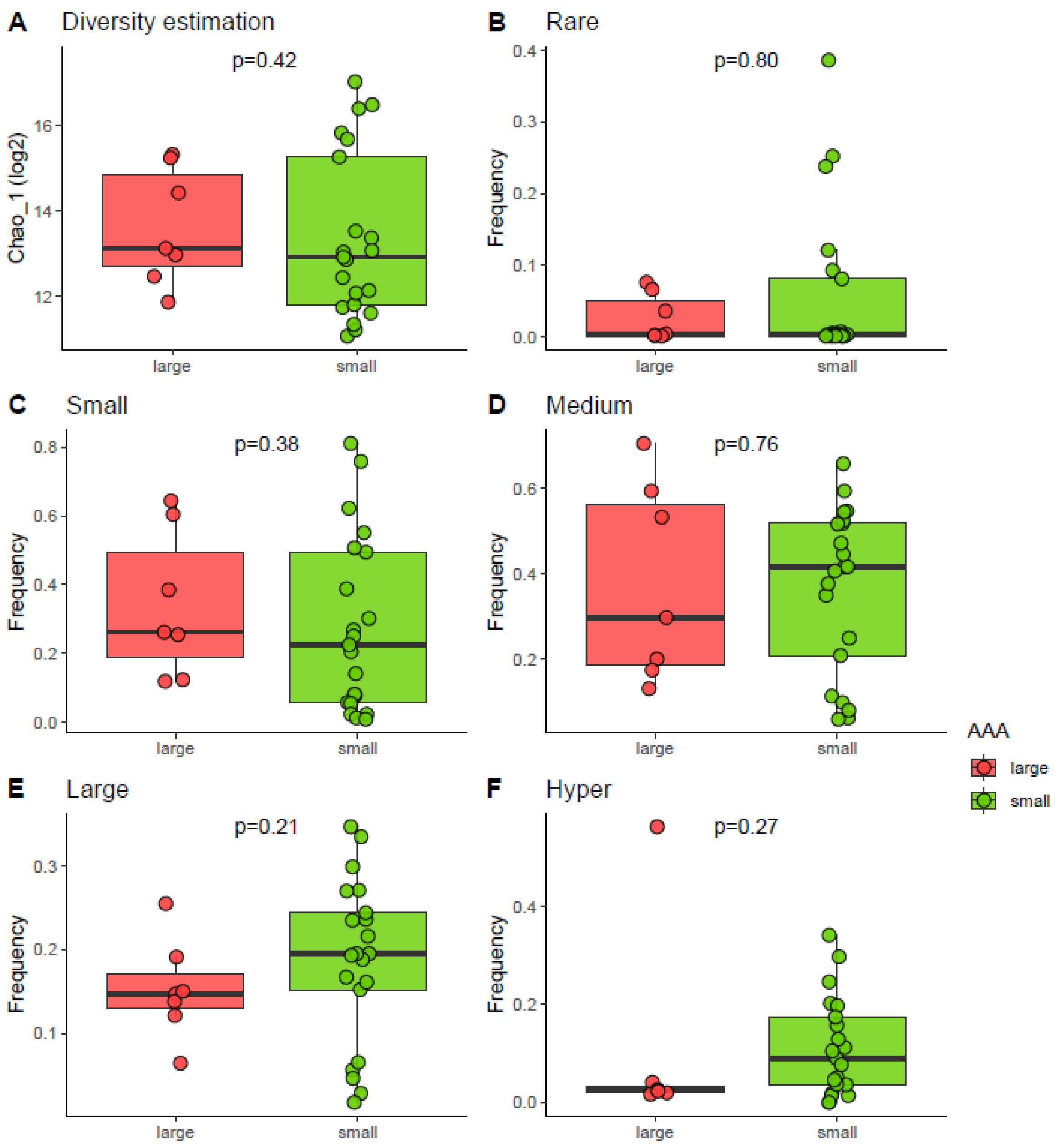


**Supplementary Figure S4. TCRβ repertoire diversity and homeostatic clonal space in large and small AAA.** Box plots of the Chao-1 ($log_2$ scaled) diversity estimate (**A**) and of the relative frequency of clonotypes (**B-F**) defined as Rare ≤0.00001; 0.00001<Small≤0.0001; 0.0001<Medium≤0.001; 0.001<Large≤0.01; and Hyperexpanded > 0.01 in large (salmon) *vs.* small (green) AAA. The two-tailed Wilcoxon test was used to assess differences between the two groups. Significance was set at P<0.05.

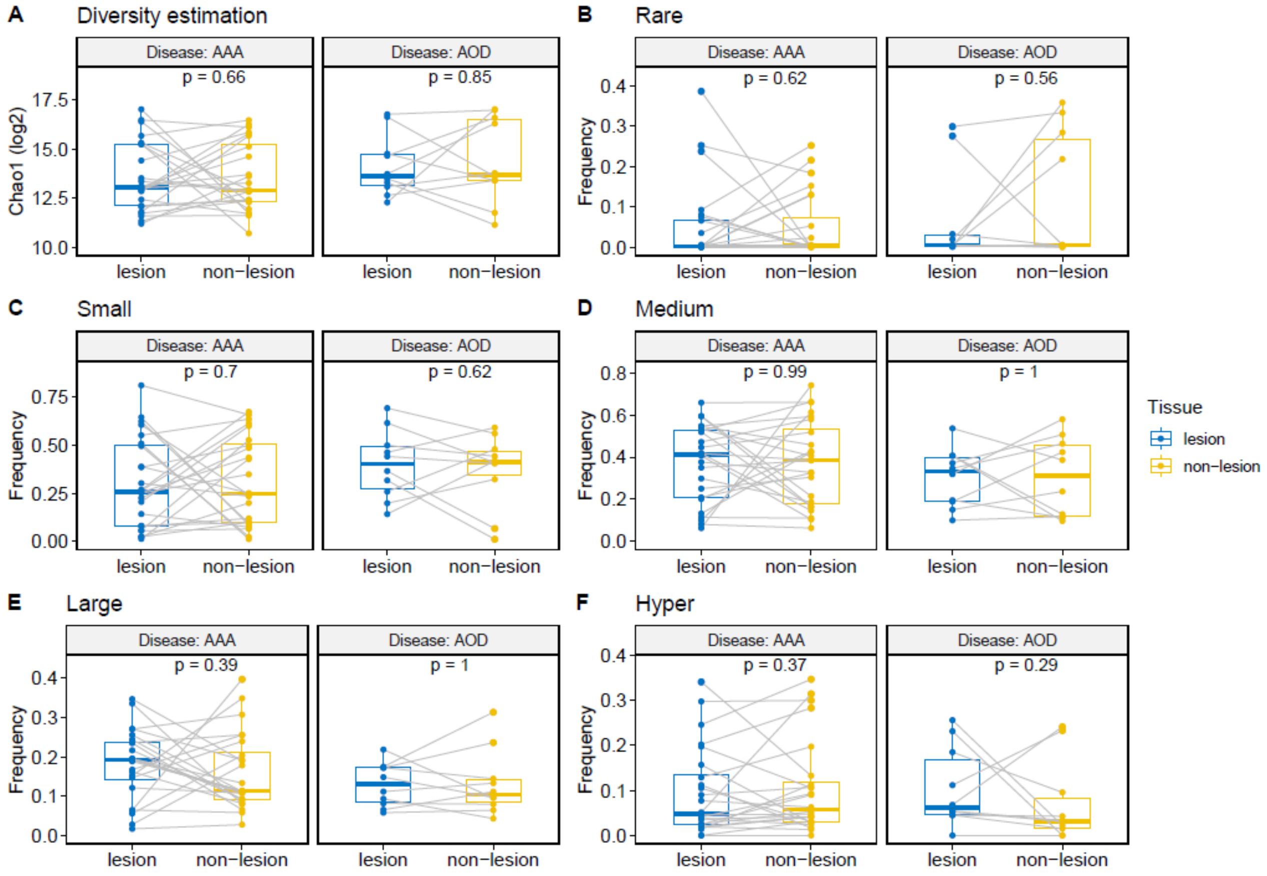


**Supplementary Figure S5. TCRβ repertoire diversity and homeostatic clonal space in PVAT of lesion *vs.* non-lesion segments within AAA and AOD.** Box plots of the Chao-1 ($log_2$ scaled) diversity estimate (**A**) and of the relative frequency of clonotypes (**B-F**) defined as Rare ≤0.00001; 0.00001<Small≤0.0001; 0.0001<Medium≤0.001; 0.001<Large≤0.01; and Hyperexpanded > 0.01 in PVAT corresponding to the lesion (blue) *vs.* non-lesion (yellow) sites for AAA and AOD paired samples. The paired Wilcoxon test was used to assess differences between the two groups. Significance was set at P<0.05.

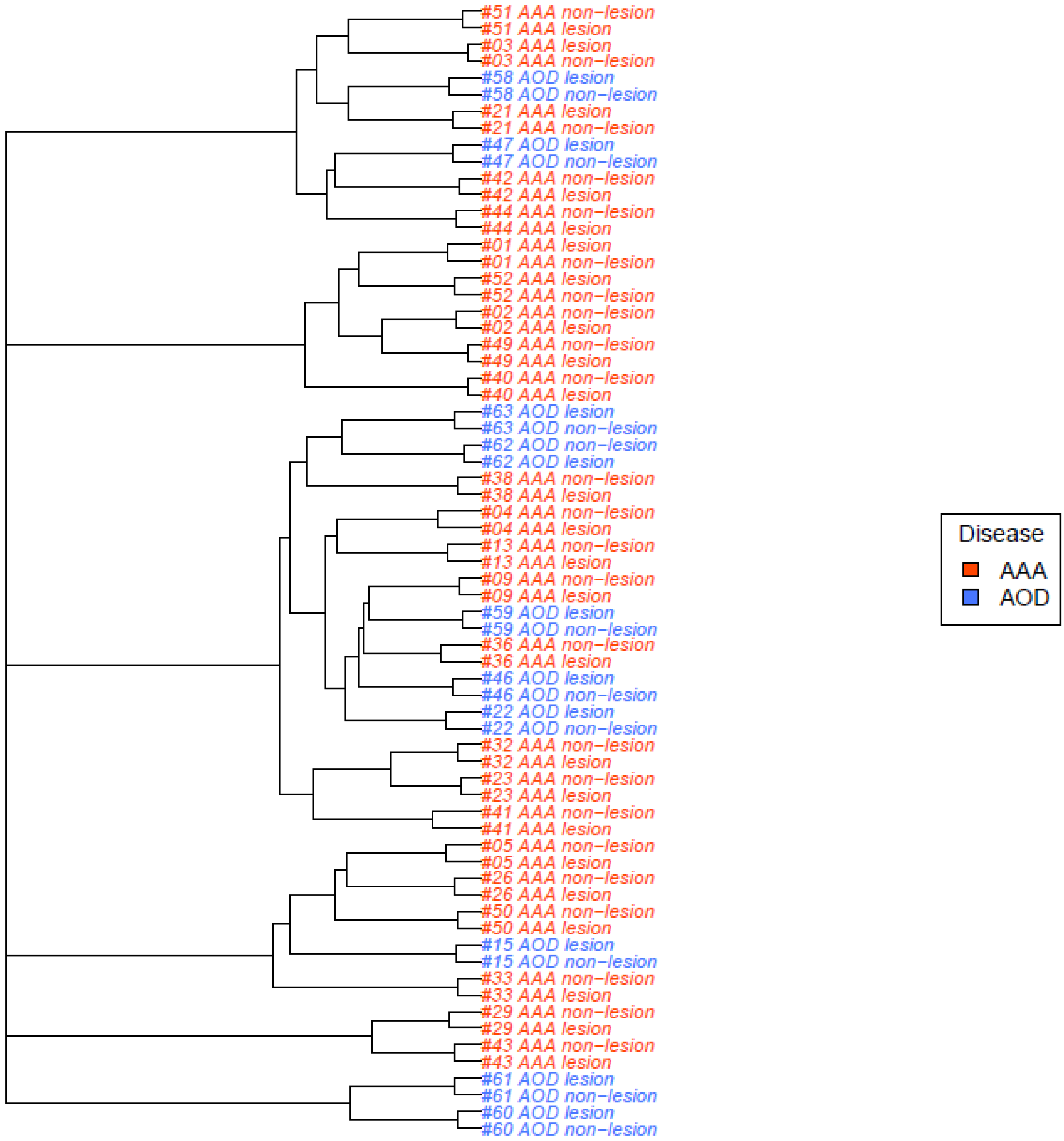


**Supplementary Figure S6. Clustering analysis of paired AAA and AOD samples.** Hierarchical clustering of the all-versus-all pairwise overlap measure (clonotype-wise sum of geometric mean frequencies, *i.e.* F2) shows that samples are clearly grouped on a patient basis, independent of disease (red=AAA; blue=AOD), suggesting that the private nature of the TCRβ repertoire drives clustering despite other phenotypic characteristics.

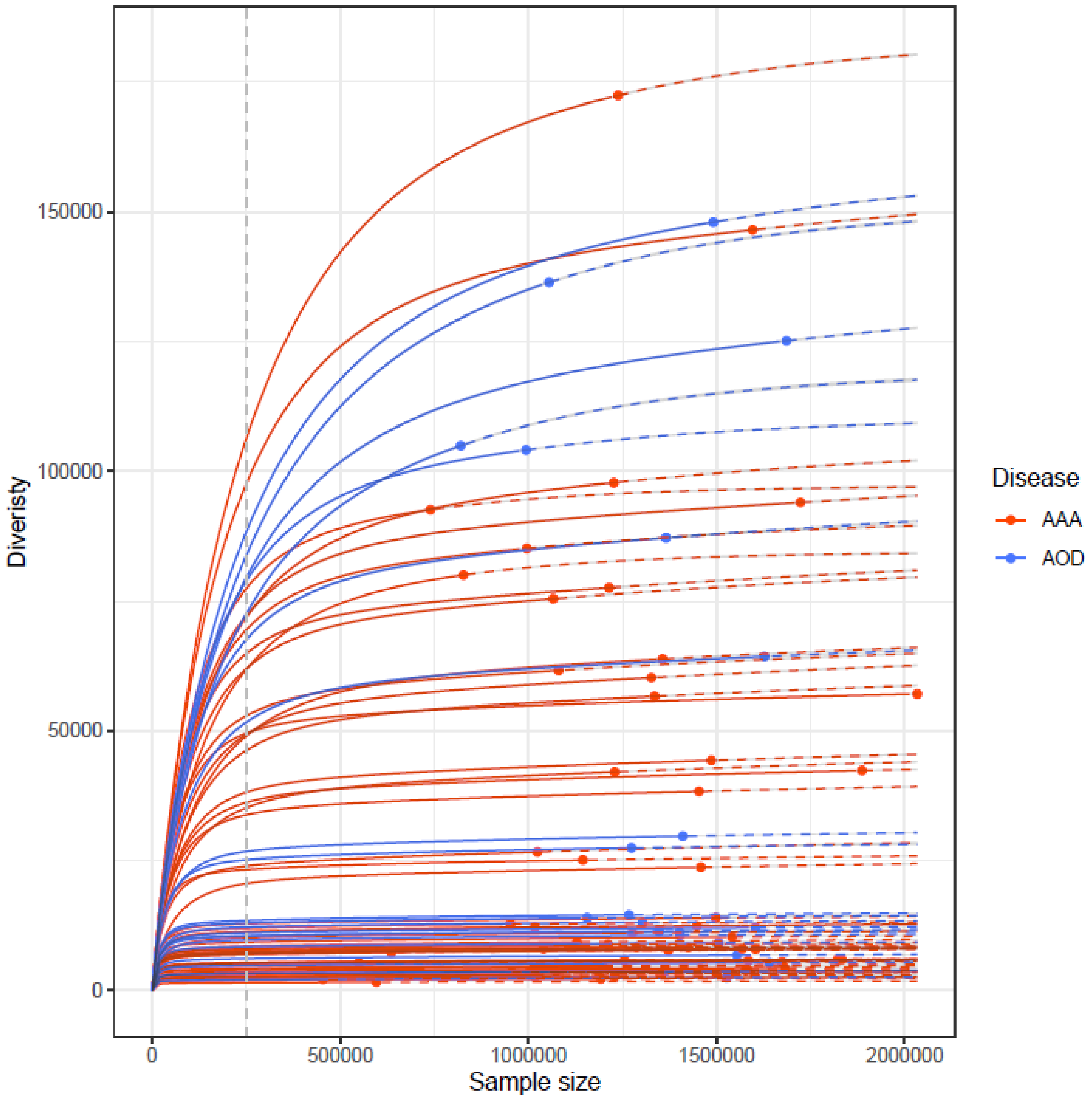


**Supplementary Figure S7. Rarefaction curves.** The graph shows the relationship between sample diversity (number of clonotypes) and sample size (reads) on raw data. The curves are interpolated (solid lines) from 0 to the current sample size and then extrapolated (dashed lines) to the size of the largest sample. Points mark the exact sample size and diversity. The dashed vertical grey line intersects all curves at the chosen threshold of 250000 reads for down-sampling and shows that most samples at this threshold are at (or close to) the plateau for diversity while remaining below their current sample size, which would reduce chip sequencing batch effects. The rarefaction curve was performed on all sequenced samples, including technical replicates. Red and blue lines indicate AAA and AOD samples respectively.

| | large (N=7) | small (N=21) | Tot AAA (N=28) | P-Val |
|---|---|---|---|---|
| **Demographics** | | | | |
| **Age, y** | | | | |
| Mean ± SD | 65.6 ± 6.65 | 68.9 ± 7.21 | 68.0 ± 7.10 | 0.29 |
| Median [Q1, Q3] | 68.0 [61.0, 70.5] | 70.0 [62.0, 73.0] | 68.5 [62.0, 72.3] | |
| **AAA max diameter, mm** | | | | |
| Mean ± SD | 69.9 ± 9.45 | 47.9 ± 2.66 | 53.4 ± 10.89 | **0.0001** |
| Median [Q1, Q3] | 69.0 [65.0, 74.6] | 48.0 [46.1, 49.5] | 48.9 [46.3, 54.4] | |
| **BMI, kg/m2** | | | | |
| Mean ± SD | 24.8 ± 1.92 | 26.4 ± 2.69 | 26.0 ± 2.59 | 0.14 |
| Median [Q1, Q3] | 24.8 [23.1, 26.4] | 26.3 [24.9, 27.8] | 25.8 [24.5, 27.3] | |
| Missing | 1 (14.3%) | 1 (4.8%) | 2 (7.1%) | |
| **CHD** | 3 (42.9%) | 9 (42.9%) | 12 (42.9%) | 1 |
| **CeVD** | 2 (28.6%) | 4 (19.0%) | 6 (21.4%) | 0.62 |
| **Hyperuricemia** | 1 (14.3%) | 0 (0%) | 1 (3.6%) | 0.22 |
| Missing | 1 (14.3%) | 0 (0%) | 1 (3.6%) | |
| **T2DM** | 1 (14.3%) | 1 (4.8%) | 2 (7.1%) | 0.40 |
| Missing | 1 (14.3%) | 0 (0%) | 1 (3.6%) | |
| **CKD** | 1 (14.3%) | 1 (4.8%) | 2 (7.1%) | 0.40 |
| Missing | 1 (14.3%) | 0 (0%) | 1 (3.6%) | |
| **Risk factors** | | | | |
| **Smokers** | 2 (28.6%) | 6 (28.6%) | 8 (28.6%) | 1 |
| Missing | 1 (14.3%) | 0 (0%) | 1 (3.6%) | |
| **Past Smokers** | 4 (57.1%) | 14 (66.7%) | 18 (64.3%) | 1 |
| Missing | 1 (14.3%) | 0 (0%) | 1 (3.6%) | |
| **Hypertension** | 4 (57.1%) | 11 (52.4%) | 15 (53.6%) | 0.66 |
| Missing | 1 (14.3%) | 0 (0%) | 1 (3.6%) | |
| **Dyslipidemia** | 4 (57.1%) | 13 (61.9%) | 17 (60.7%) | 1 |
| Missing | 1 (14.3%) | 0 (0%) | 1 (3.6%) | |
| **Laboratory tests** | | | | |
| **Total cholesterol, mg/dL** | | | | |
| Mean ± SD | 200 ± 30.6 | 192 ± 33.6 | 194 ± 32.5 | 0.56 |
| Median [Q1, Q3] | 196 [178, 221] | 198 [169, 211] | 197 [171, 211] | |
| **HDLc, mg/dL** | | | | |
| Mean ± SD | 34.0 ± 6.81 | 43.2 ± 10.0 | 40.9 ± 10.0 | **0.02** |
| Median [Q1, Q3] | 37.0 [28.5, 38.0] | 43.0 [37.0, 50.0] | 41.5 [35.3, 47.3] | |
| **Triglycerides, mg/dL** | | | | |
| Mean ± SD | 183 ± 89.5 | 115 ± 52.4 | 132 ± 68.8 | 0.06 |
| Median [Q1, Q3] | 176 [110, 242] | 104 [73.0, 136] | 120 [84.8, 148] | |
| **Medications** | | | | |
| **Antiaggregants** | 6 (85.7%) | 17 (81.0%) | 23 (82.1%) | 1 |
| **Diuretics** | 1 (14.3%) | 4 (19.0%) | 5 (17.9%) | 1 |
| **ACE inhibitors** | 1 (14.3%) | 4 (19.0%) | 5 (17.9%) | 1 |
| **Statins** | 2 (28.6%) | 9 (42.9%) | 11 (39.3%) | 0.67 |
| **Beta blockers** | 4 (57.1%) | 5 (23.8%) | 9 (32.1%) | 0.17 |
| **Calcium blockers** | 2 (28.6%) | 5 (23.8%) | 7 (25.0%) | 1 |
| **Proton pump inhibitors** | 3 (42.9%) | 9 (42.9%) | 12 (42.9%) | 1 |
| **Oral hypoglycemics** | 1 (14.3%) | 1 (4.8%) | 2 (7.1%) | 0.44 |
| **ARBs** | 2 (28.6%) | 3 (14.3%) | 5 (17.9%) | 0.57 |

**Supplementary Table S1. AAA subgroups patient clinical characteristics.** Categorical variables are shown as counts (n) and percentage (%); quantitative demographic variables are expressed both as mean ± standard deviation (SD) and as median and interquartile range [Q1, Q3]. Medications are summarized by drug class. Comparisons of large vs. small AAA for categorical variables were made by Fisher's exact test, when appropriate. Continuous variables were compared by 2-tailed t-test or Wilcoxon test for non-normally distributed variables. Statistical significance for comparison: $P<0.05$. AAA indicates abdominal aortic aneurysm; ACE, angiotensin-converting-enzyme; AOD, Aortic Occlusive Disease; ARBs, angiotensin II receptor blockers; BMI, body mass index; CeVD, cerebrovascular disease; CHD, coronary heart disease; CKD, chronic kidney disease; HDL-c, high-density lipoprotein cholesterol.
